# Proximity labeling and orthogonal nanobody pulldown (ID-oPD) approaches to map the spinophilin interactome uncover a putative role for spinophilin in protein homeostasis

**DOI:** 10.1101/2025.01.23.634546

**Authors:** Emily T. Claeboe, Keyana L. Blake, Nikhil R. Shah, Cameron W. Morris, Basant Hens, Brady K. Atwood, Sabrina Absalon, Amber L. Mosley, Emma H. Doud, Anthony J. Baucum

## Abstract

Spinophilin is a dendritic spine enriched scaffolding and protein phosphatase 1 targeting protein. To detail spinophilin interacting proteins, we created an Ultra-ID and ALFA-tagged spinophilin encoding construct that permits proximity labeling and orthogonal nanobody pulldown (ID-oPD) of spinophilin-associated protein complexes in heterologous cells. We identified 614 specific, and 312 specific and selective, spinophilin interacting proteins in HEK293 cells and validated a subset of these using orthogonal approaches. Many of these proteins are involved in mRNA processing and translation. In the brain, we determined that spinophilin mRNA is highly neuropil localized and that spinophilin may normally function to limit its own expression but promote the expression of other PSD-associated proteins. Overall, our use of an ID-oPD approach uncovers a novel putative role for spinophilin in mRNA translation and synaptic protein expression specifically within dendritic spines.

## INTRODUCTION

Classically, serine/threonine protein phosphatase catalytic subunits dephosphorylate a great number of substrates, suggesting promiscuous activity. However, in practice, phosphatases, such as protein phosphatase 1 (PP1), complex with various regulatory and/or inhibitory protein partners to enhance specificity of protein targeting^1–4^. There are over 200 known PP1 targeting proteins. While targeting of PP1 regulatory subunits may hold therapeutic promise^5,6^, we must first detail specific substrates for selected PP1 holoenzymes. Given a dearth of effective and specific drugs to treat neuropsychiatric and neurological disorders, detailing how neuronal PP1 regulatory proteins associate with specific PP1 substrates may have therapeutic implications.

Spinophilin (*PPP1R9B*) is the most abundant neuronal, postsynaptic density (PSD)-enriched PP1 targeting protein. Loss of spinophilin basally increases locomotor output and decreases anxiety-like behaviors in rodents but also limits neuroadaptations underlying psychostimulant-induced locomotor sensitization, rotarod motor learning, and excessive repetitive motor output associated with mouse models for excessive grooming^7–12^. Additionally, spinophilin expression and/or protein interactions within the striatum are increased by psychostimulants, which increase striatal dopamine, and are decreased by a 6-hydroxydopamine mouse lesion model associated with loss of nigrostriatal dopamine neurons^13–16^. Using co-immunoprecipitation and proteomic approaches, we have identified multiple classes of spinophilin interacting proteins, including cytoskeletal, ribosomal, PSD, and heat shock proteins isolated from neuronal and non-neuronal cells^16–18^. Importantly, we have found that these interactions are regulated by dopamine depletion, psychostimulant treatment, and obesity^15,16,18^. Additionally, we have found cell type-specific regulation of striatal behaviors and metabolic parameters in conditional spinophilin knockout (KO) mice^10,18^. However, the use of co-immunoprecipitation affinity purification approaches to tease apart the valid interactors from the non-specific or high abundance proteins, termed the CRAPome, can be challenging^19^. Therefore, validated, orthogonal purification approaches are needed to enhance the sensitivity, specificity, and rigor of previously defined interactomes.

We transfected cells with a spinophilin protein that had an ALFA-UltraID protein Cre-dependently fused to its C-terminus. We identified 614 spinophilin interacting proteins that were “specific,” meaning they were detected in the presence of our spinophilin construct when it was co-transfected with Cre-recombinase but not when the spinophilin construct was transfected alone. However, we found that many of the high abundance, specific spinophilin interactors were not “selective” as they were detected in control experiments that used an EGFP-tagged UltraID. Therefore, there is high specificity, but low selectivity, for overexpression proximity labeling studies. When subtracting out the non-selective interacting proteins, we detected 312 spinophilin interacting proteins that were both specific and selective.

To detail functional regulation of protein expression by spinophilin-dependent targeting of PP1, we overexpressed a wildtype (WT) spinophilin or a PP1 binding-deficient spinophilin mutant (F451A)^9,20^. We found that basal expression of F451A mutant spinophilin was greater than WT spinophilin. Conversely, we detected less expression of the spinophilin interacting proteins, such as ribosomal subunits, in the mutant compared to wildtype spinophilin. This may suggest spinophilin normally promotes ribosomal protein expression and subsequent function (e.g. mRNA translation). Importantly, we found that spinophilin mRNA has punctate expression in the neuropil of the striatum whereas mRNA for the spinophilin homolog, neurabin, is localized to the nuclear/perinuclear area. Additionally, we found that spinophilin limits its own expression *in vivo* as we detected minimal cell type-specific viral transduction of spinophilin in wildtype, but not cell type-specific spinophilin KO, mice. In contrast to limiting its own expression, loss of spinophilin decreased the expression of PSD proteins that are known to have highly neuropil-translated mRNAs. Together, our data suggest a putative role for spinophilin in regulating PSD protein organization and expression, uncovering a novel potential role for spinophilin in dendritic spine protein synthesis.

## RESULTS

### Generation and validation of a Cre-dependent spinophilin-ALFA-UltraID construct

We created and verified a plasmid encoding an ALFA-tag and UltraID protein Cre-dependently appended to the C-terminus of spinophilin (**Figure 1A**). We transfected two different clones of this plasmid into HEK293 cells in the absence or presence of a plasmid encoding Cre recombinase. Lysates were precipitated with streptavidin magnetic beads or neutravidin agarose beads. ALFA-tag immunoreactivity was only present when Cre recombinase was co-transfected (**Figure 1B**). We detected ALFA-tag immunoreactivity in the pulldowns, demonstrating a biotinylation of our spinophilin protein. When samples were immunoblotted for spinophilin, two bands were present in the Cre-expressing cells. In contrast, only the lower spinophilin band was present when Cre was absent, corresponding to untagged spinophilin expression that occurs in the absence of Cre (**Figure 1B**). In addition to spinophilin, we detected PP1 in the neutravidin pulldowns only when Cre recombinase was co-transfected (**Figure 1B**). Streptavidin signal was robustly detected when the ALFA-UltraID-spinophilin was co-expressed with Cre recombinase, demonstrating a high level of specific labeling (**Figure 1B**). These data demonstrate, and validate, the generation of a Cre-dependent ALFA– and UltraID-tagged spinophilin that permits robust detection of known spinophilin interacting proteins, such as PP1.

**Figure 1.**
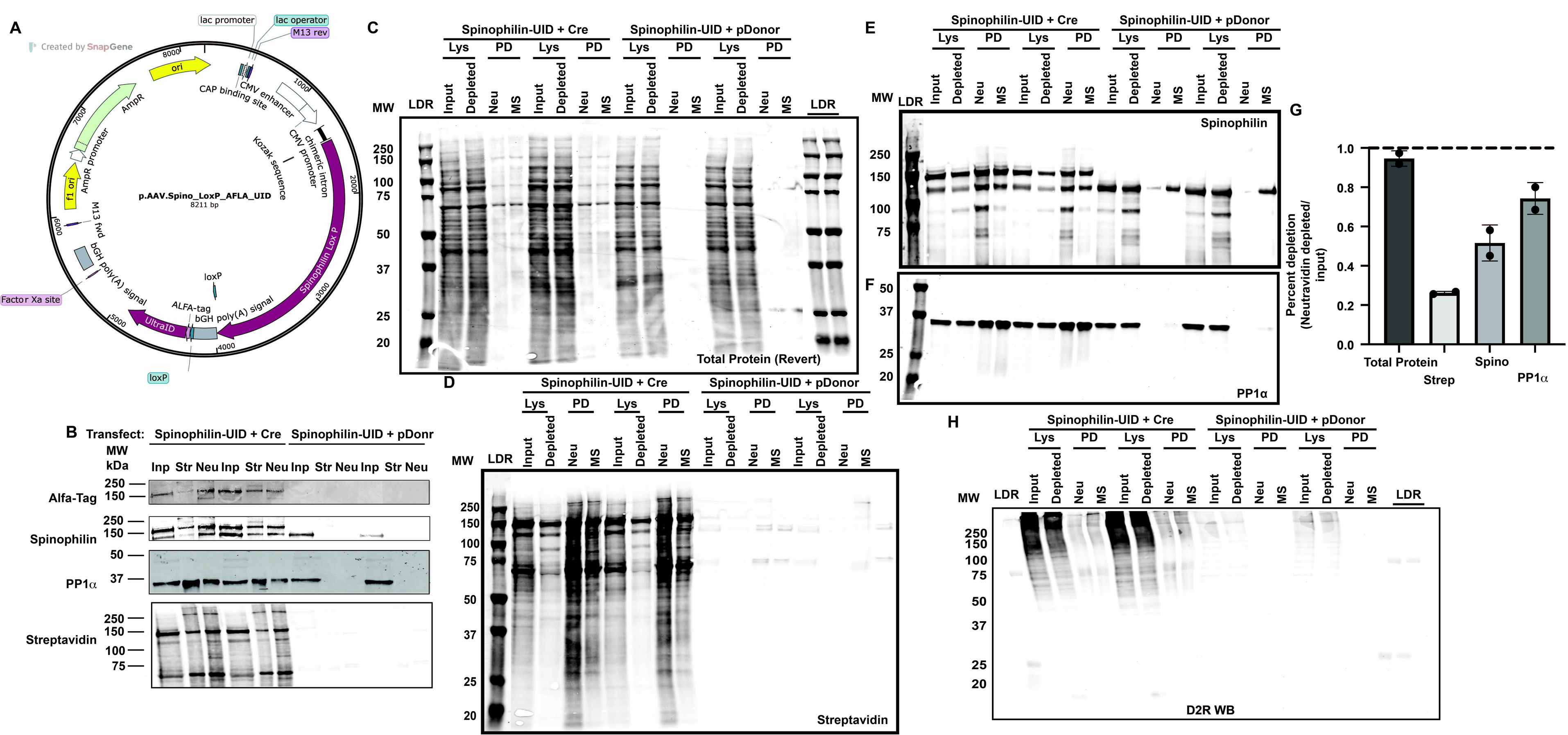
Generation and validation of a Cre-dependent spinophilin-ALFA-UltraID construct. **A.** Map of the sequence validated spinophilin-ALFA-UltraID construct created in SnapGene. **B.** HEK293 cells were transfected with the spinophilin-ALFA-UltraID construct (Spinophilin-UID) along with a construct encoding an improved Cre recombinase (Cre) or an empty pDonr221 vector (pDonr). Following transfections, cells were incubated with biotin. HEK293 lysates were then precipitated using streptavidin magnetic beads (Str) or neutravidin agarose beads (Neu). Inputs (Inp) and pulldowns were separated by SDS-PAGE and immunoblotted with an ALFA-tag (Alfa), spinophilin, or PP1α antibody or probed with an Alexa Dye 800-conjugated streptavidin protein. Blots were imaged using a LiCor Odyssey M. **C.** HEK293 cells were transfected with the spinophilin-ALFA-UltraID construct (Spinophilin-UID) along with a construct encoding an improved Cre recombinase (Cre) or an empty pDonr221 vector (pDonr). Cells were also transfected with dopamine D2 receptor. Following transfections, cells were incubated with biotin. HEK293 lysates were then precipitated using neutravidin agarose beads (Neu) or trypsin resistant streptavidin magnetic beads (MS). Inputs, lysates (Lys), or post neutravidin pulldown lysates (depleted) or Neu or MS pulldowns (PD) were separated by SDS-PAGE and probed with a revert, total protein stain (**C**), an Alexa Dye 800-conjugated streptavidin protein (**D**) or immunoblotted for spinophilin (**E**) or PP1α (**F**). **G. D.** HEK293 cells were transfected with the spinophilin-ALFA-UltraID construct (Spinophilin-UID) along with a construct encoding an improved Cre recombinase (Cre) or an empty pDonr221 vector (pDonr). Cells were also transfected with dopamine D2R. Following transfections, cells were incubated with biotin. HEK293 lysates were then precipitated using Neu or MS beads. Inputs, Lys, or depleted, or Neu or MS PD were separated by SDS-PAGE and immunoblotted with a D2R antibody. **H.** The total protein, streptavidin, spinophilin, or PP1α signal intensity in the depleted lysate was divided by the signal intensity in the input to determine the efficiency of Neu pulldown.

Biotinylated proteins were isolated using neutravidin or a trypsin resistant streptavidin that is chemically modified to limit streptavidin digestion by trypsin, thereby enhancing the signal to noise in mass spectrometry-based proteomics applications^21^. We detected equal total protein in both the input lysate and the neutravidin depleted lysate lanes, suggesting that the biotinylated proteins that are precipitated make up a small fraction of the total protein complement (**Figures 1 C, G**). We observed robust total protein (Revert stain) and streptavidin staining in the neutravidin and the trypsin-resistant streptavidin protein pulldown lanes when Cre recombinase was co-expressed, but minimal total protein and streptavidin fluorescence in the absence of Cre recombinase (**Figure 1C**). We observed ∼80% decrease in streptavidin labeling (**Figures 1 D, G**), ∼50% loss in spinophilin immunoreactivity (**Figures 1 E, G**), and ∼25% decrease in PP1 immunoreactivity (**Figures 1F, G**), in the depleted supernatant following neutravidin pulldown. We observed spinophilin and PP1 in both neutravidin and streptavidin pulldown conditions. Additionally, there was some low level of non-specific binding of the MS-streptavidin to overexpressed spinophilin (**Figure 1E**). However, even with pulldown of spinophilin, co-precipitating PP1 was minimally detected in the neutravidin pulldowns of non Cre-recombinase expressing cells (**Figure 1F**). Cells used for neutravidin or MS streptavidin pulldown were also co-transfected with GFP-tagged D2R. We immunoblotted the streptavidin pulldowns using a D2R antibody and detected a smeary signal in the immunoprecipitates when Cre-recombinase was expressed, but not in the absence of Cre-recombinase (**Figure 1H**). Overexpression of GPCRs, like the D2R, commonly leads to a smeary banding pattern.

### Proteomics analysis of spinophilin interactome

While we can detect spinophilin association with GPCRs such as mGluR5 and D2R by immunoblotting^10^, we have never detected GPCR proteins in spinophilin immunoprecipitates from brain lysates using mass spectrometry^16–18^. Therefore, a portion of the streptavidin pulldowns from the pulldowns using trypsin-resistant streptavidin were reserved and submitted for tryptic digestion, mass spectrometry, and a search against the human database for peptide spectral matching. We detected 10 peptide spectral matches (PSMs) matching D2R and 8 matching GFP (termed a contaminant as present in the contaminant database) (**Table 1A**).

**Table 1.**
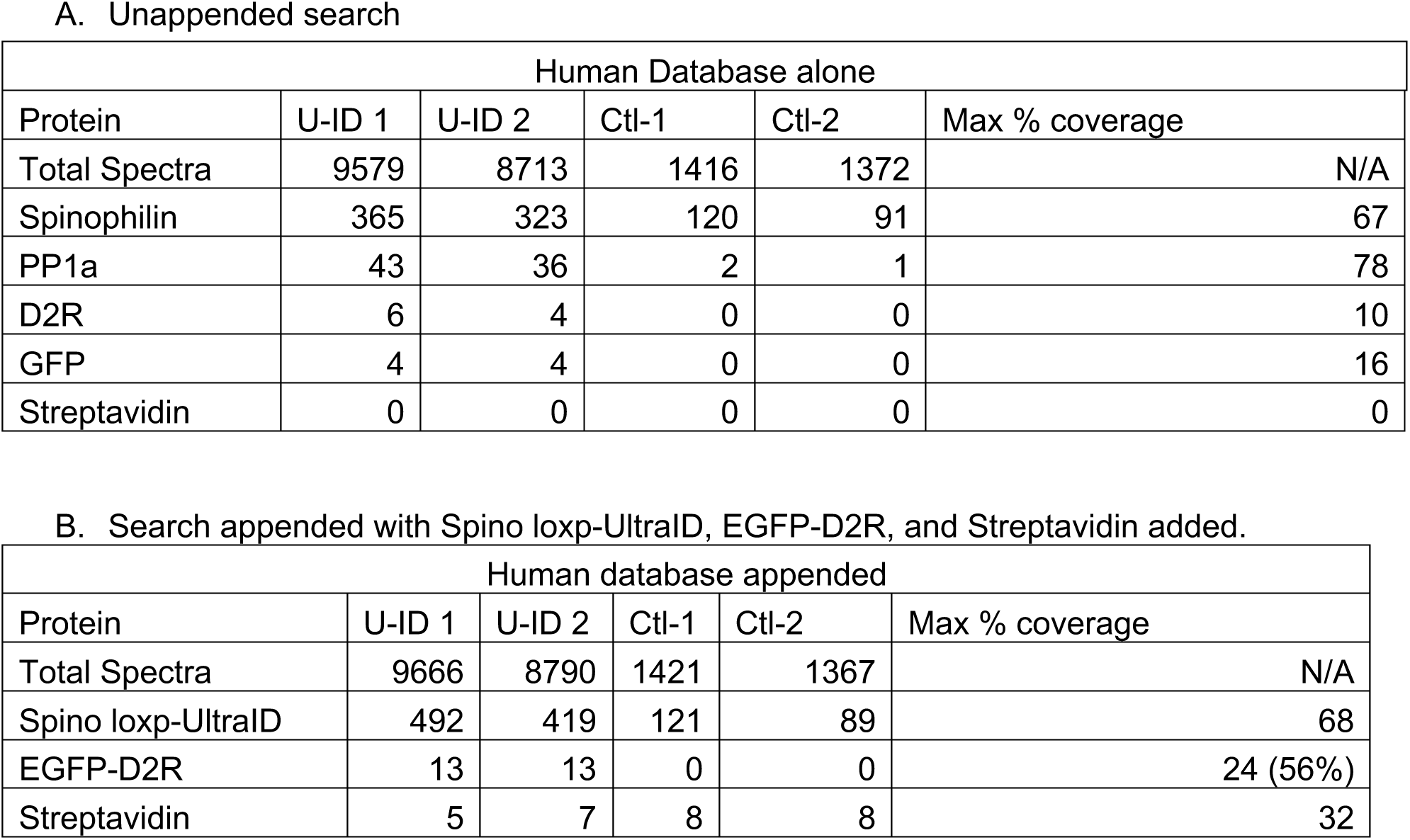
– Peptide spectral matches (PSMs) detected from proteomics analysis of trypsin resistant streptavidin pulldowns.

We re-searched the human database with the sequences for GFP-D2R, Spino-loxp-ALFA-UltraID, and streptavidin appended to the database. We detected more PSMs to the EGFP-D2R appended sequence than the total PSMs matching D2R or GFP in the un-appended search (**Table 1**). We observed 56% coverage of the GFP sequence but only 24% coverage of the total sequence (**Table 1B**). We observed a greater number of PSMs matching the Spino-loxp-ALFA-UltraID when these sequences were appended for the database search only in the samples expressing this construct, demonstrating specific expression in the presence of Cre recombinase (**Table 1B**). Consistent with specific biotinylation and pulldown, there was a 6.6-fold greater number of PSMs from the Ultra-ID expressing cells compared with the control cells (**Table 1A**). The number of peptide spectral matches (PSMs) matching spinophilin was ∼3.2-fold greater in the Cre-recombinase expressing cells with an average of 344 PSMs in the experimental condition and 106 in the control condition (**Table 1A**). Spinophilin was unexpectedly present in the control condition, potentially due to high expression and background binding of the overexpressed spinophilin (without UltraID tag) to the trypsin-resistant streptavidin beads. However, we detected minimal interacting proteins in the negative control. For example, PP1 isoforms were either highly enriched (19-26-fold) or selectively detected in the spinophilin-Ultra-ID expressing samples compared to spinophilin-alone overexpression (**Tables 1A, S1**). We also only detected an average of 7 PSMs per sample matching streptavidin suggesting the trypsin-resistant streptavidin is lowly abundant, limiting concern of signal suppression (**Table 1B**).

In addition to GFP-tagged D2R, we detected 806 proteins with at least 2 PSMs in our spinophilin proximity labeling proteomics studies (**Table S1A**). Of the 806 total proteins, 743 were either specifically enriched by >3.2-fold in spinophilin-Ultra-ID vs control cells (**Table S1B**) or were not common contaminants. A total of 619 proteins that had a total of at least 4 PSMs across the two samples were detected (**Table S1B – bolded proteins**).

### Spinophilin pathway enrichment

We analyzed the specific spinophilin interactome which contained 615 proteins that were present in the string database (**Table S1C**, String-db) ^22–25^. The top 10 “component” gene ontology (GO) pathways are enriched in ribosomal and focal adhesion proteins (**Figure 2A**). The top 10 “molecular function” GO pathways are enriched in proteins associated with binding and translation functions (**Figure 2B**). The top 10 “biological process” GO pathways are enriched in proteins associated with translation and metabolism (**Figure 2C**). The top 10 Kyoto Encyclopedia of Genes and Genome (KEGG) pathways are enriched in ribosomal proteins, RNA transport, and neurological disorders (**Figure 2D**). Many of the proteins associated with the different pathways have high overlap. Using the String-db to evaluate the 81 proteins that match to the “Cytoplasmic Translation” in the “Biological Process” enrichment term, we identified multiple ribosomal proteins and translation initiation proteins (**Figure 2E**). The full complement of pathways for the GO and KEGG enrichments are given in **Tables S2A-S2D**. In addition to the above pathways, we also evaluated the WikiPathways, a newer, open-source, pathway database tool^26^. There was high overlap between WikiPathways and the GO and KEGG analyses, including ribosomal proteins and translation. In addition to these pathways, Alzheimer’s disease pathways and Vascular Endothelial growth factor (VEGFA-VEGFR2) signaling were in the top 10 pathways (**Table S2E**).

**Figure 2.**
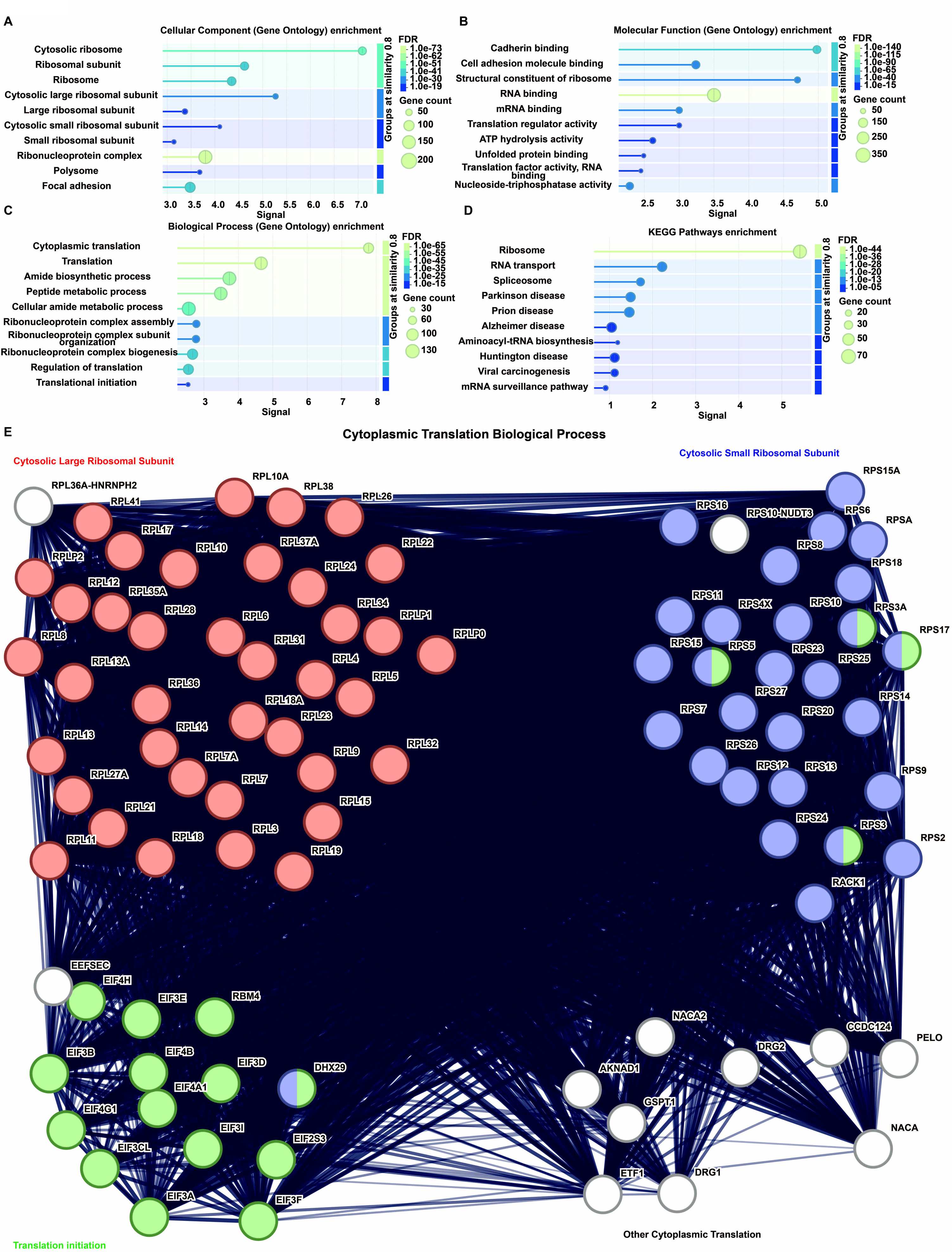
Connectivity and function of specific spinophilin interacting proteins. The specific spinophilin interacting proteins based on peptide spectral matches (PSMs) were input into the string database (www.string-db.org). The database matched 615 total proteins. The false discovery rate (FDR), signal, and gene count for top 10 Gene Ontology (GO) enrichment terms matching **A.** “cellular component”, **B.** “molecular function, **C.** “biological process” as well as the **D.** Kyoto encyclopedia of genes and genomes (KEGG) enrichment terms are shown. **E.** The proteins matching the “cytoplasmic translation” biological process and their string interactions are plotted. Within this enrichment, proteins matching the “cytosolic large ribosomal subunit”, “cytosolic small ribosomal subunit”, and “translation initiation” were enriched. Network edges are confidence based with the line thickness indicating how well the data support the connection.

### Validation and quantitation of spinophilin interactions by proximity biotinylation and ALFA-tag nanobody co-immunoprecipitation

We validated the proteomics studies by immunoblotting neutravidin and trypsin resistant streptavidin pulldowns. We detected spinophilin and PP1α similarly in both pulldowns. We also validated Bip (Gpr78), 40S ribosomal protein S3, and beta-tubulin as specific interactors in both pulldowns (**Figure 3A**). We also detected spinophilin and PP1α in the ALFA-tag nanobody pulldown (**Figure 3B**). We detected both Bip and beta tubulin in our neutravidin and ALFA nanobody pulldowns (**Figure 3B**). We detected 40S ribosomal protein S3 in the neutravidin pulldowns; however, this protein was very faint in the ALFA-tag pulldowns (**Figure 3B**). Importantly, we did not detect spinophilin or the interacting proteins in non-transfected cells.

**Figure 3.**
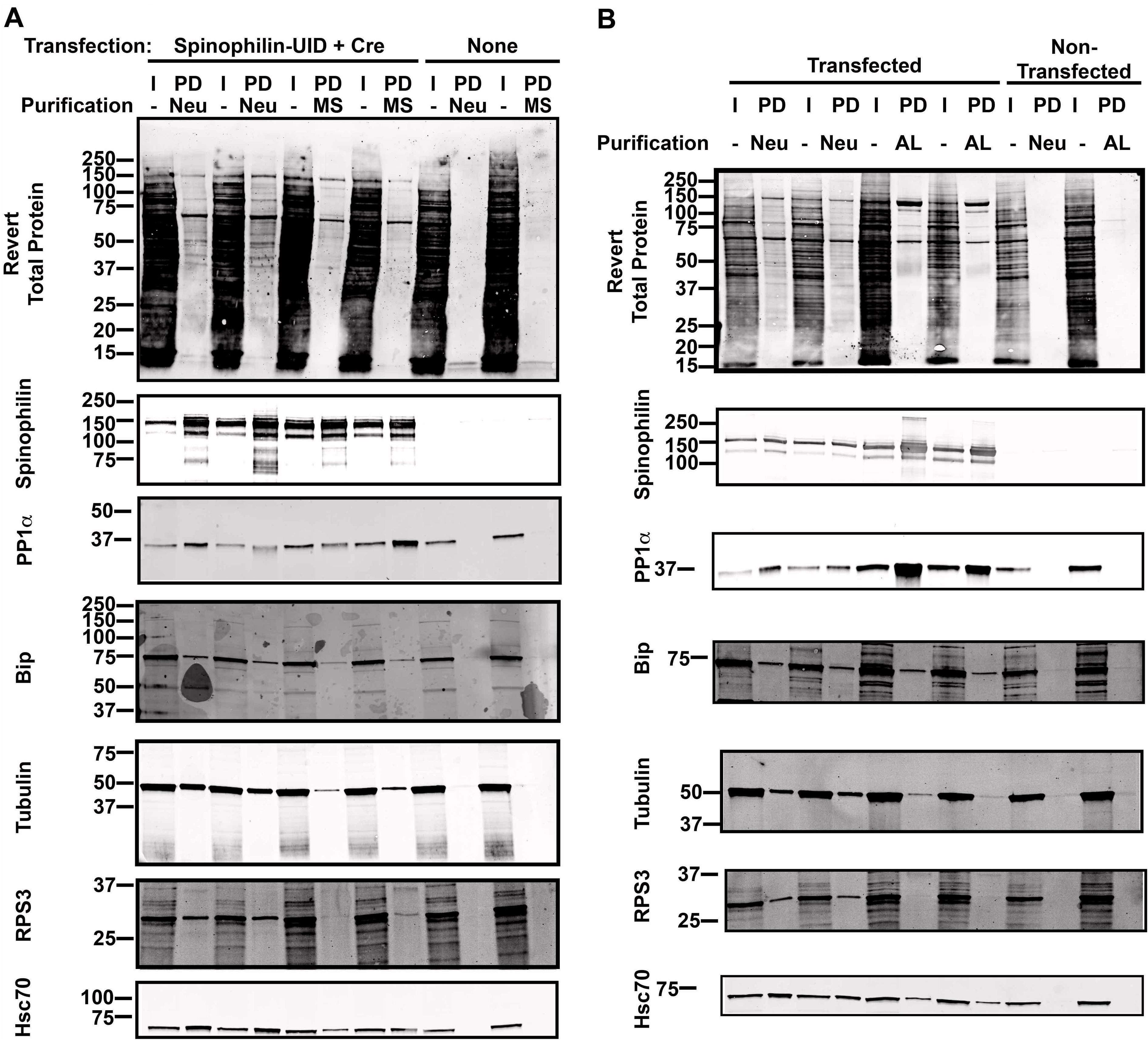
Comparison of neutravidin pulldown and ALFA-tag nanobody immunoprecipitation to trypsin-resistant streptavidin beads. HEK293 cells were transfected with the spinophilin-ALFA-UltraID construct (Spinophilin-UID) along with a construct encoding an improved Cre recombinase (Cre) or were non-transfected. **A.** Following transfections, cells were incubated with biotin. HEK293 lysates were then precipitated using neutravidin agarose beads (Neu) or trypsin resistant streptavidin magnetic beads (MS). Inputs (I) and pulldowns (PD) were separated by SDS-PAGE and imaged for total protein with Revert stain and subsequently immunoblotted for spinophilin, PP1α, Bip, Beta-tubulin (tubulin), Rps3, or Hsc70. Blots were imaged using a LiCor Odyssey M. **B.** Following transfections, cells were incubated with biotin. HEK293 lysates were then precipitated using neutravidin agarose beads (Neu) or an ALFA-tag nanobody fused to agarose beads (AL). Inputs (I) and pulldowns (PD) were separated by SDS-PAGE and imaged for total protein with Revert stain and subsequently immunoblotted for spinophilin, PP1α, Bip, Beta-tubulin (tubulin), Rps3, or Hsc70. Blots were imaged using a LiCor Odyssey M.

### Selectivity of spinophilin-dependent interactome mapping

Together, we have found that the use of our ID-oPD approach permits identification and validation of specific spinophilin protein interactions. However, many of these proteins are associated with mRNA translation and may non-selectively interact with *any* transfected protein. To detail the selective spinophilin interactome, we overexpressed EGFP that had UltraID directly fused to its C-terminus and analyzed the protein interactions using proteomics approaches as above (**Table S3**). The construct used had the synapsin promoter to permit neuronal expression, but previous studies have shown that this promoter also drives protein expression in HEK293 cells and consistent with this, we detected the expressed construct by proteomics^27^. Importantly, when analyzing the interactome, many of the pathways associated with EGFP interactome overlap with the spinophilin interactome (**Figure S1**). These data suggest that while proteins are “specific,” as defined above, many are not “selective” as they associated with UltraID fused to a different protein. However, 313 specific proteins were detected in the spinophilin UltraID studies that had at least 4 PSMs and that were not observed in the EGFP pulldowns (**Table S4**). Importantly, we did not detect proteins such as PP1 in the EGFP UltraID pulldown. The 312 (out of the 313 proteins detected) proteins that were present in the string-db database were searched and pathway analysis was performed. When the EGFP-dependent non-selective interactors were removed, many of the same pathways, such as “RNA binding proteins” and “translation”, were still associated with the spinophilin interactome (**Figure 4A-D**). Moreover, “cellular metabolic process” term was now in the top 10 biological processes, suggesting proteins within this area are both specific and selective spinophilin interactors. When evaluating the specific and selective protein interactors that map to “cellular metabolic process” we identified proteins within “primary metabolic process”, “RNA binding”, and “translation” subcategories (**Figure 4E**). The full enrichments are given in **Table S5A-S5E**.

**Figure 4.**
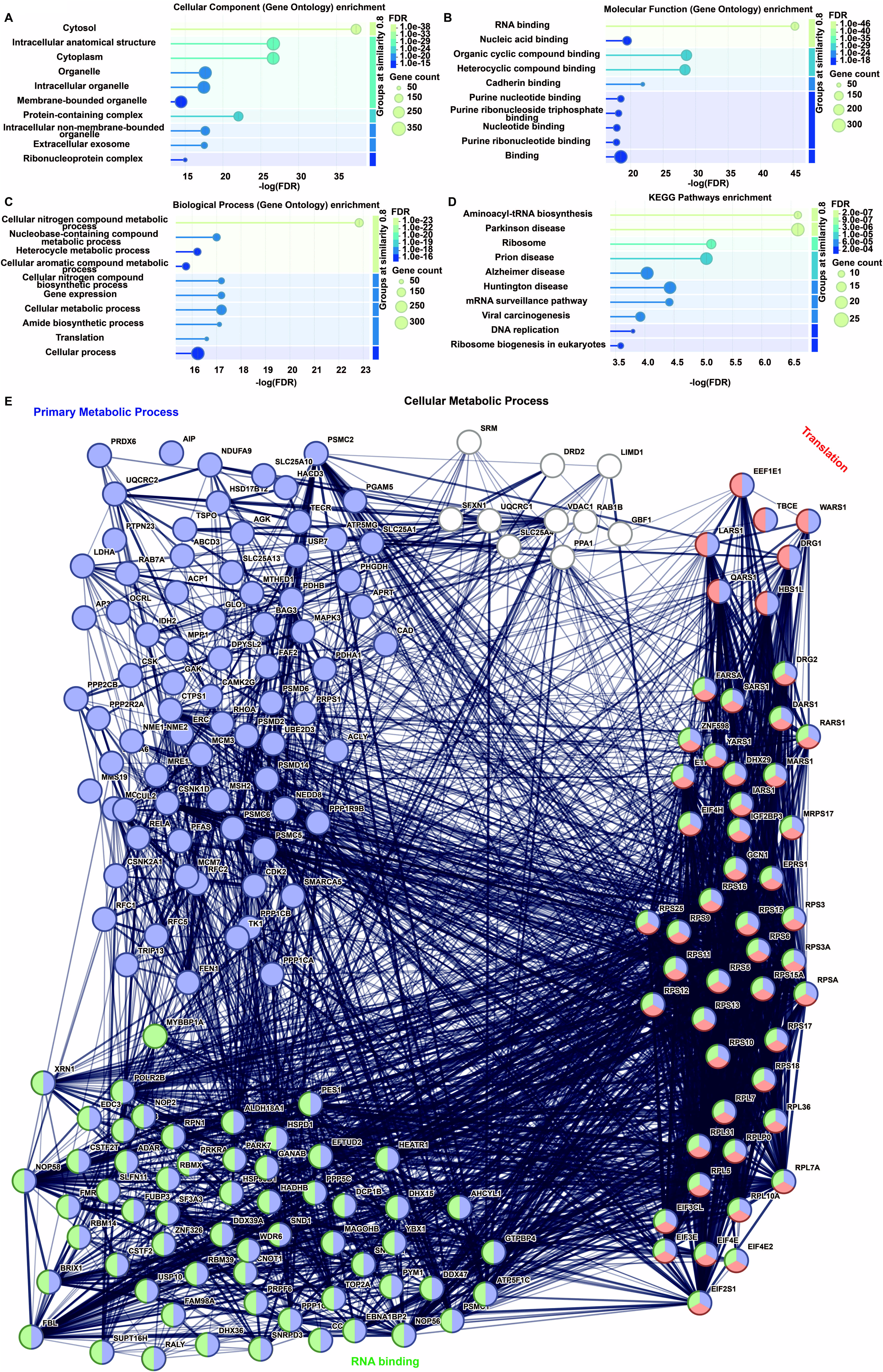
Connectivity and function of specific and selective spinophilin interacting proteins. The specific and selective spinophilin interacting proteins based on peptide spectral matches (PSMs) were input into the string database (www.string-db.org). The database matched 312 total proteins. The false discovery rate (FDR), signal, and gene count for top 10 Gene Ontology (GO) enrichment terms matching **A.** “cellular component”, **B.** “molecular function, **C.** “biological process” as well as the **D.** Kyoto encyclopedia of genes and genomes (KEGG) enrichment terms are shown. **E.** The proteins matching the “cellular metabolic process” within the biological GO enrichment and their string interactions are plotted. Within this enrichment, proteins matching the “primary metabolic process” and “translation” were enriched. Network edges are confidence based with the line thickness indicating how well the data support the connection.

### Spinophilin-dependent targeting of PP1 limits spinophilin expression but stabilizes expression of proteins involved in protein homeostasis

The use of EGFP-UltraID tagged controls suggests that these highly specific approaches may lack selectivity, at least in the context of overexpression studies. However, even if non-selective, spinophilin may be modulating the expression and/or function of these interacting proteins. To determine how spinophilin may regulate the expression of cellular proteins in a PP1-dependent manner, we overexpressed wildtype or a PP1 binding-deficient (F451A) mutant spinophilin in HEK293 cells. We performed tandem mass tag (TMT) quantitative proteomics on the cell lysates isolated from a single well of a 6-well plate. We detected a total of ∼660 proteins that were quantified across all samples (**Table S6A; Figure 5A**). Of these, 146 proteins had a log_2_ fold-change of ±0.5 or greater increase or decrease in expression in the mutant compared with wildtype expressing cells (**Figure 5A**). Consistent with our interactome data, we detected enrichment of ribosomal proteins, cellular metabolic processes, and cytoplasmic translation (**Figure 5B-E**). When plotting the interactome for the detected proteins matching the cellular metabolic process, as in our selective interactome, we identify multiple proteins matching “primary metabolic process” and “translation” (**Figure 5F**). Interestingly, 136 of the 146 proteins that had altered expression demonstrated a decreased expression in the mutant cells, suggesting an overall positive regulation of protein expression by wildtype spinophilin. Of the 136 decreased proteins, 62 were identified in the spinophilin-UltraID pulldown. In contrast to most of the altered proteins having decreased expression, we found that mutant spinophilin had a ∼2-fold greater expression than wildtype spinophilin when normalized to the total protein abundance (**Figure 5A**). Those proteins that had both a trend towards significance (p<0.1) and a log_2_ fold-change > ±0.5 detected in the mutant compared to wildtype overexpressing cells are shown in **Figure S2**. These proteins were enriched in “RNA binding proteins” (p=0.0043) within the “Molecular Function” GO enrichment and the “metabolism of proteins” (p=3.70e-05) within the “Reactome Pathway” enrichment. Overall, spinophilin-dependent targeting of PP1 increases protein expression of multiple spinophilin interacting proteins associated with metabolism, protein homeostasis, and mRNA function whereas spinophilin binding to PP1 seems to negatively regulate its own expression.

**Figure 5.**
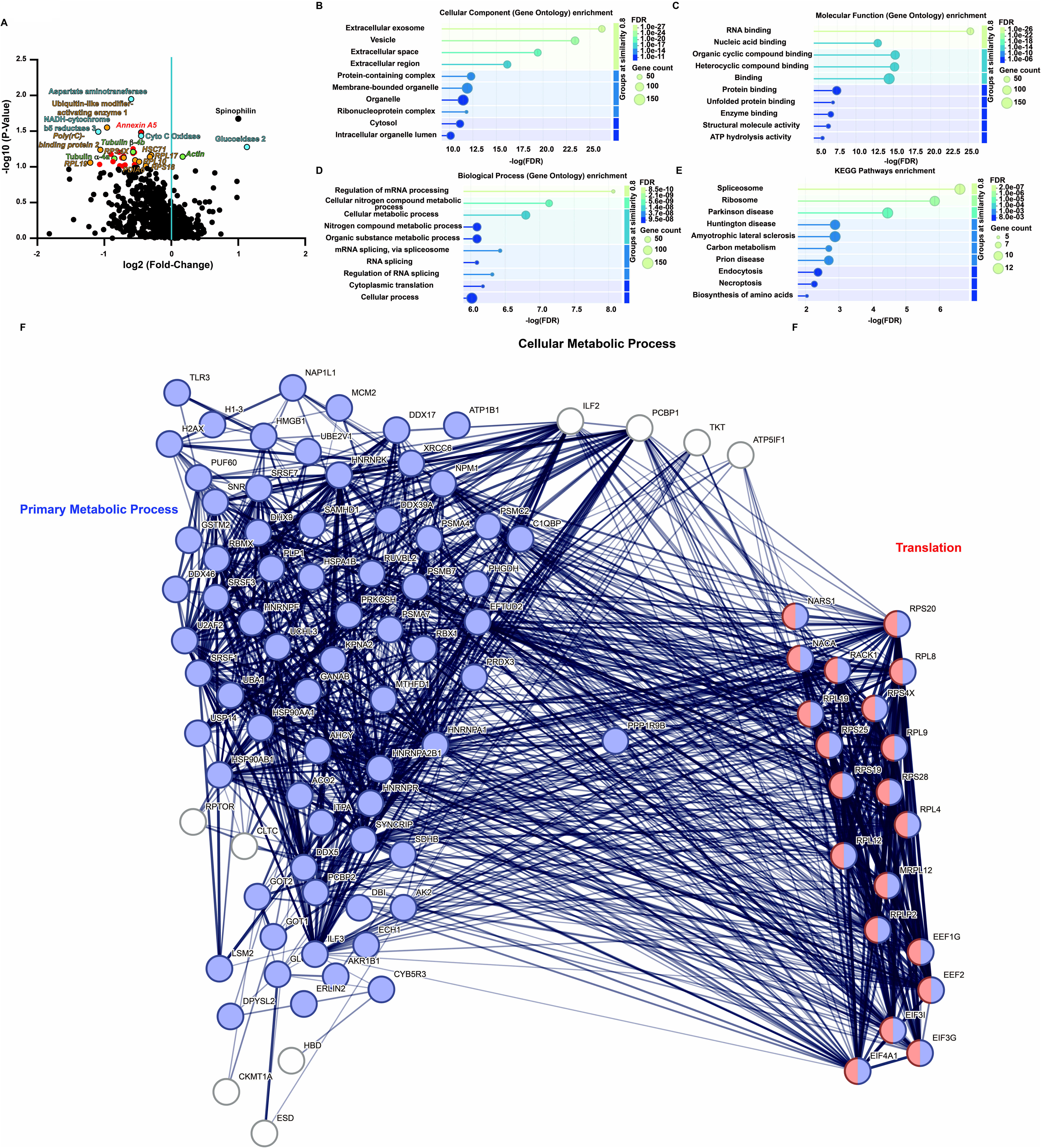
Connectivity and function of proteins differentially expressed in spinophilin F451A mutant compared to wildtype overexpressing cells. Proteins with a log_2_ fold-change of ± 0.5 in the F451A overexpressing HEK293 cells compared with the wildtype spinophilin overexpressing cells were input into the string database (www.string-db.org). The database matched 146 total proteins. The false discovery rate (FDR), signal, and gene count for top 10 Gene Ontology (GO) enrichment terms matching **A.** “cellular component”, **B.** “molecular function, **A**. “biological process” as well as the **D.** Kyoto encyclopedia of genes and genomes (KEGG) enrichment terms are shown. **E.** The proteins matching the “cellular metabolic process” within the biological GO enrichment and their string interactions are plotted. Within this enrichment, proteins matching the “primary metabolic process” and “translation” were enriched. Network edges are confidence based with the line thickness indicating how well the data support the connection. **F.** F451A mutant spinophilin had significantly greater abundance of the Tandem mass tags compared with wildtype spinophilin when overexpressed in heterologous cells.

While our studies were focused on total protein abundance, we did detect 22 phosphorylated peptides matching 19 phosphorylation sites on 16 proteins. The only significantly different phosphorylated peptide matched spinophilin (**Table S6**). However, this peptide phosphorylation was increased as was the total protein expression and therefore, there is no difference in phosphorylation of the peptide when normalized to total protein (**Figure S3**). Moreover, when normalizing those peptides that came from proteins that had multiple non-phosphorylated peptides detected, none of these detected phosphorylation sites were significantly impacted by overexpression of F451A mutant, compared with wildtype, spinophilin (**Figure S3**). Some of the peptides detected had only one phosphorylated PSM or a single major contributing PSM. We compared the non-normalized PSM abundance values from these phosphopeptides and again there was no significantly different phosphorylation (**Figure S4**). While not significant, there were two phosphorylated peptides that are involved in RNA function that trended towards a differential phosphorylation: acid nuclear phosphoprotein 32 family member B (ANP32B) and RNA-binding protein 39 (RBM39).

### *Ppp1R9b* (spinophilin) mRNA is diffusely expressed and spinophilin limits its own expression

Our HEK cell studies demonstrate that spinophilin interacts with multiple ribosomal subunits and mRNA translation pathways. Moreover, spinophilin protein is highly enriched in dendritic spines^28,29^. Previous, unbiased studies suggested that spinophilin mRNA was also enriched in hippocampal neuropil^30,31^. Therefore, to understand how spinophilin may modulate these pathways *in vivo* and to validate these unbiased studies, we evaluated the localization of spinophilin (*Ppp1r9b*) mRNA using RNAScope. Overall, *Ppp1r9b* mRNA is expressed throughout the brain (**Figure 6A**). There are both overlapping and unique enrichments of the mRNA of the spinophilin homolog, neurabin (*Ppp1r9a*). Intriguingly, at higher magnifications, spinophilin mRNA staining is localized as discrete puncta, suggesting a neuropil localization throughout the striatum (**Figure 6B**). This contrasts with the spinophilin homolog, *Ppp1r9a* mRNA, which is nuclear/perinuclear.

**Figure 6.**
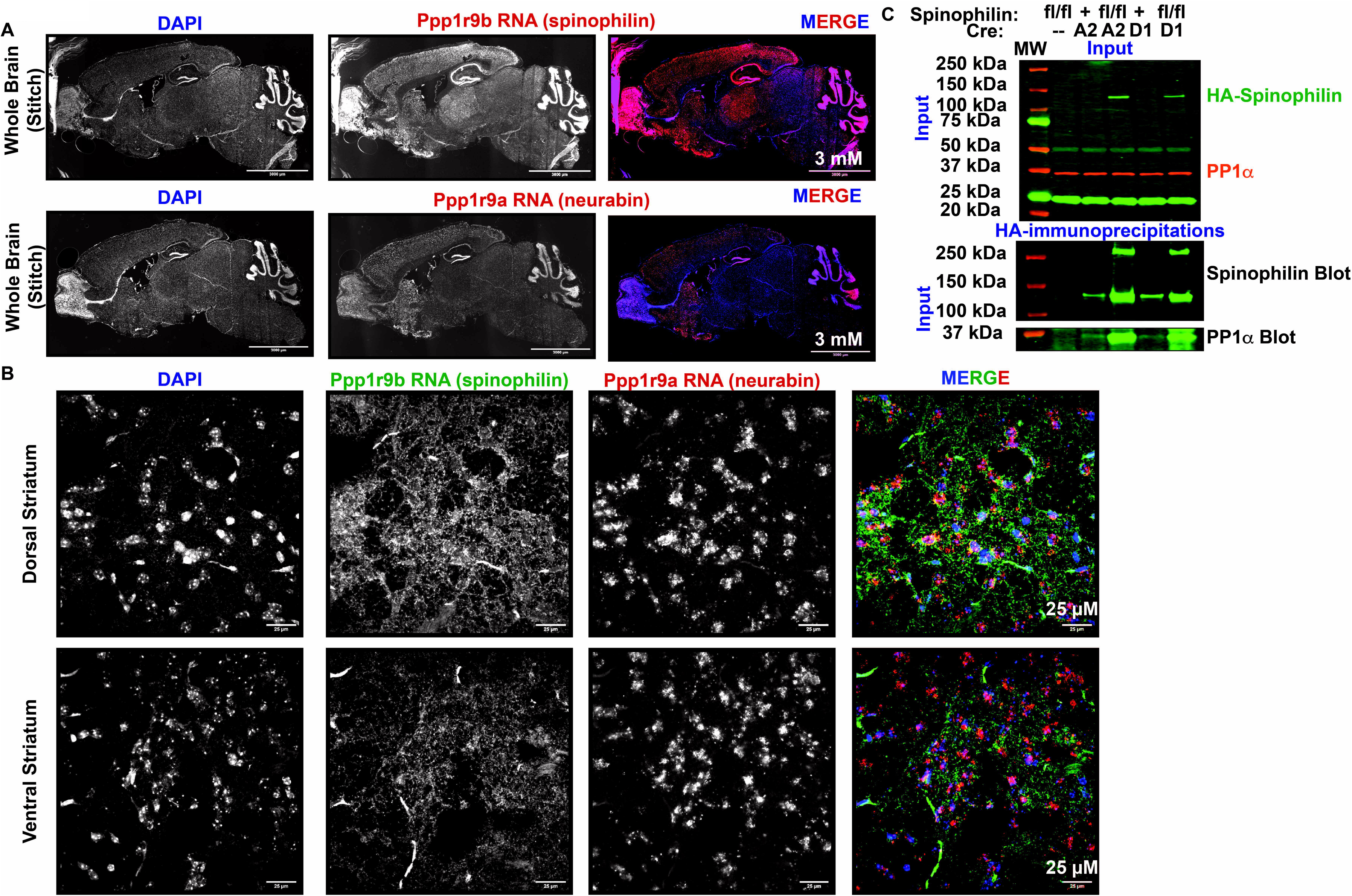
Unique subcellular distribution of spinophilin (Ppp1r9b) and neurabin (Ppp1r9a) mRNA and a role for spinophilin in regulating its own expression. RNAScope analysis of wildtype mouse brains at 4X stitched together (**A**) and 63X magnification (**B**). **C.** Spinophilin^fl/fl^/A2A Control mice (Spino^fl/fl^ or mice expressing AdorA2A-Cre (A2A) or Drd1-Cre (D1)) or mice with loss of spinophilin in iMSNs (Spinophilin^fl/fl^/A2A) or dMSNs (Spinophilin^fl/fl^/D1) were transduced with a virus encoding a Cre-dependent, HA-tagged spinophilin. Total lysates (Input) were immunoblotted for HA epitope tag and PP1α and HA-immunoprecipitates were immunoblotted for spinophilin and PP1α.

We created a virus that encoded a Cre-dependent HA-tagged spinophilin to permit overexpression of a tagged form of spinophilin within direct pathway medium spiny neurons (dMSNs), using a Drd1-Cre mouse line, or indirect pathway medium spiny neurons (iMSNs), using an AdorA2A mouse line. There was very little expression of the HA-spinophilin in spinophilin control Cre-expressing mice, even though this protein was expressed using a robust, cytomegalovirus (CMV) promoter (**Figure 6C**). This may be due to the unique mRNA localization of *Ppp1r9a* being required for expression of the protein. However, viral transduction of mice lacking endogenous spinophilin in dMSNs or iMSNs had robust expression of HA-spinophilin (**Figure 6C**), suggesting that the viral construct can be expressed, but the endogenous spinophilin is somehow acting to limit its own expression *in vivo*.

### Spinophilin promotes expression of specific synaptic proteins

While spinophilin limits its own expression, our HEK293 cell studies suggest that wildtype spinophilin promotes expression of multiple proteins when compared with overexpression of F451A spinophilin. To determine how loss of spinophilin impacts striatal synaptic protein expression, we used a fractionation protocol to isolate synaptoneurosome and PSD fractions from striatum of spinophilin wildtype and KO mice^32^. Cytosolic proteins (GAPDH, PP1α, and PP1ψ1) were enriched in the total homogenate whereas PSD proteins (PSD95 and GluN2B) were enriched in the PSDs (**Figures 7A, B**). We evaluated the expression of known PSD scaffolding and receptor proteins, including PSD95, GluN2B, and GluA2 as well as two PP1 isoforms (PP1α and PP1ψ1) in the synaptoneurosome and PSD fractions of spinophilin wildtype and global spinophilin KO mice. When normalized to the amount of protein detected in the total homogenate, there was a trend for a decreased protein expression of PP1α in the synaptoneurosome fraction in the KO compared with wildtype mice (**Figure 7C**). In contrast all proteins that were detected in the PSD fraction had decreased expression in the KO compared with the wildtype mice in the PSD fraction when protein abundance was normalized to the protein’s abundance in the total homogenate fraction (**Figure 7D**). Consistent with decreases of all proteins, there was a trend (p=0.05) or significant decreases in the total protein levels in the synaptoneurosome and PSD fractions, respectively (**Figure 7E**). When individual proteins within each fraction were normalized to the total protein stain, we observed an *increased* expression of PSD95 in the total homogenate, but no difference in the other two fractions. Conversely, when normalized to total protein, we observed a trend or significant decreases in PP1 levels across all fractions. (**Figures 7F-H**). Therefore, spinophilin seems to promote the expression, but not targeting, of PSD-associated proteins whereas it may promote both the expression and targeting of PP1 isoforms to the PSD.

**Figure 7.**
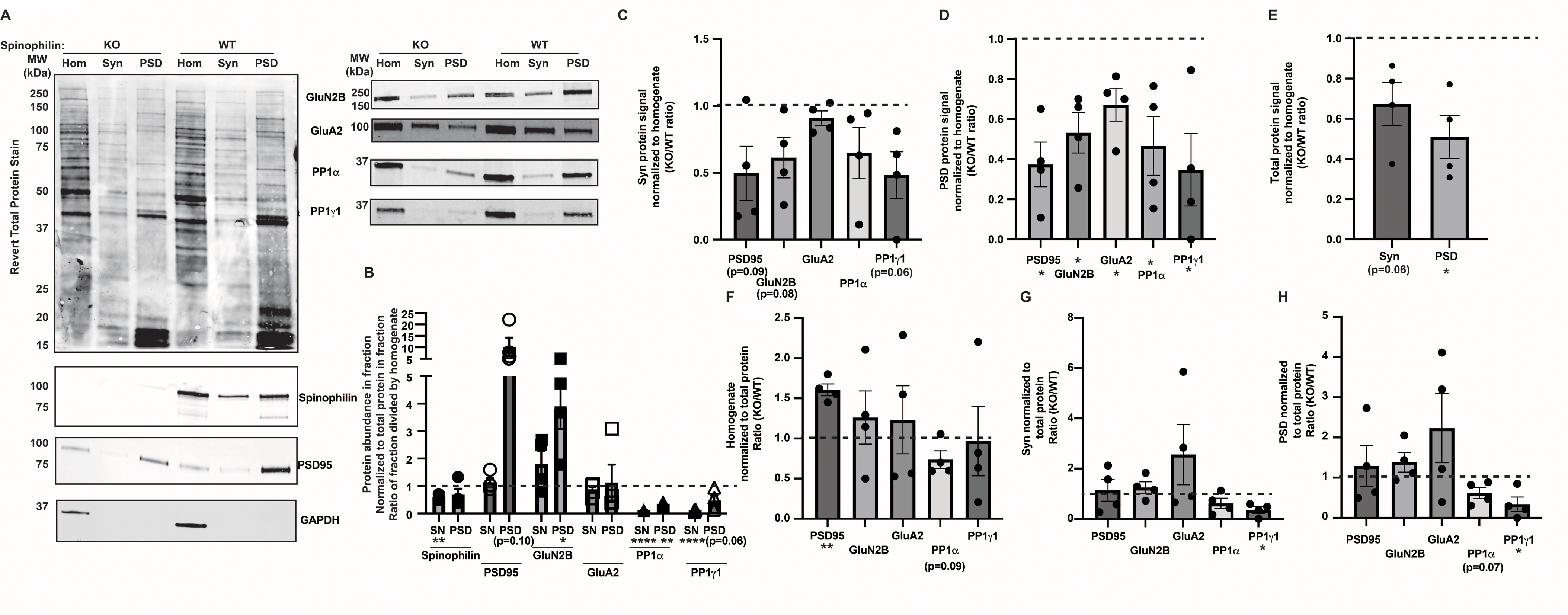
Spinophilin functions to stabilize PSD proteins. Striata from wildtype (WT) or spinophilin knockout (KO) mice were homogenized and biochemically fractionated into synaptoneurosome (Syn), or postsynaptic density (PSD) fractions. (**A**) Total homogenate (Hom), syn, and PSD fractions were separated by SDS page and stained with Revert total protein stain or immunoblotted for spinophilin, PSD95, GAPDH, GluN2B, GluA2, PP1α, and PP1ψ1. (**B**) The fluorescence intensity of the specific protein in the homogenate, Syn, or PSD fraction from spinophilin WT mice was normalized to the total protein abundance within the corresponding fraction. A ratio was generated by dividing the syn or PSD fraction values by the homogenate values. (**C, D**) The fluorescence intensity of the specific protein in the Syn (**C**) or PSD (**D**) fraction isolated from spinophilin WT or KO mice was divided by the fluorescence intensity of the protein within the corresponding homogenate fraction. A ratio was generated by dividing the KO value by the WT value. (**E**) The fluorescence intensity of the total protein stain in the Syn or PSD fraction isolated from spinophilin WT or KO mice was divided by the fluorescence intensity of the total protein stain within the corresponding homogenate fraction. A ratio was generated by dividing the KO value by the WT value. (**F, G, H**) The fluorescence intensity of the specific protein in the homogenate (**F**), Syn (**G**), or PSD (**H**) fraction isolated from spinophilin WT or KO mice was divided by the fluorescence intensity of the total protein stain within the corresponding fraction. A ratio was generated by dividing the KO value by the WT value. See Supplemental Western Figure for all full blots used for quantitation. See Table S8 for all statistics analyses.

## DISCUSSION

### ID-oPD as a specific and sensitive approach to identify the spinophilin interactome

We generated a novel ALFA and UltraID-tagged spinophilin. Unlike HA and Myc tags, the ALFA epitope tag is not expressed in any known mammalian system^33^, does not require folding for antibody detection, and can be readily immunoprecipitated using specific camelid nanobodies^33^. UltraID is one of the newest generations of BioID *in vivo* proximity labeling proteins. It is one of the smallest, most robust, and most specific proximity ligases generated to date, having higher thermal stability than miniTurbo along with less background labeling and a smaller size than TurboID^34^. Additionally, to limit signal suppression by high abundance, non-specific proteins, we utilized a trypsin resistant streptavidin to isolate biotinylated proteins^21^. This approach led to limited detection of streptavidin PSMs, with an average of 7 PSMs per pulldown across the 4 conditions. This is ∼1.5% of the number of PSMs of the target spinophilin UltraID protein. Thus use of this streptavidin would limit signal suppression and enhance detection of low abundance proteins, ameliorating concerns of streptavidin contamination for proteomics approaches^35^. In addition to low streptavidin background, we found that there was a high level of enrichment of biotinylated proteins in the experimental compared with the control sample. Specifically, we detected an average of 9,146 PSMs across the 2 experimental conditions compared with 1,394 PSMs across the 2 control conditions. Also, many of the proteins that were similarly detected in the experimental and control conditions were known endogenously biotinylated proteins such as carboxylases and histones. These proteins that directly bind the streptavidin beads can be subtracted out by use of a non-UltraID protein expressing control^36,37^.

We utilized targeted immunoblotting to validate a subset of the mass spectrometry results. Moreover, to permit orthogonal validation of interacting proteins, we used an ALFA-tag pulldown. The recently described ALFA-tag permits isolation of the linear epitope and is a sequence that is not expressed in the mammalian system^33^. Using nanobody pulldowns we observed robust blotting for spinophilin and PP1α. While we detected Bip, Tubulin, RPS3, and HSC70 in the ALFA-tag nanobody pulldowns, the signal for RPS3 was very faint. These data suggest that proximity labeling and nanobody pulldowns permit orthogonal validation of our proteomics data; however streptavidin-based pulldowns may be more robust and permit better quantification of protein interactome changes.

### Proximity labeling identifies specific interactors associated with chaperone function

Multiple studies show that spinophilin interacts with, and regulates the function of PP1, cytoskeletal proteins, and postsynaptic density-enriched proteins^9,10,16,17,20,38–40^. We and others have also identified other classes of proteins, such as ribosomal proteins, chaperone proteins, and mitochondrial proteins that are more broadly distributed outside the brain^16–18,41^. However, the specificity of these interactions is unclear as many of these protein classes are common contaminants in affinity purification studies and are part of the contaminant repository for affinity purification mass spectrometry data, or CRAPome^19^.

We detected “unfolded protein binding” as one of the top 10 molecular functions. Two of the most abundant spinophilin interacting proteins were heat shock proteins; heat shock 70 kDa protein 1b (Hsc70; *HSPA1B*) and heat shock cognate 71 kDa protein (Hsc71; *HSPA8*). These two proteins have high levels of non-specific association as reported in the CRAPome^19^. Based on PSMs, there was a ∼10-fold and ∼50-fold enrichment in the experimental conditions for Hsc70 and Hsc71, respectively. This contrasts with our previous co-immunoprecipitation studies where we detected an equal number of PSMs or 4-fold more Hsc71 when performing immunoprecipitations with a mouse or goat spinophilin antibody, respectively, from wildtype compared with spinophilin KO mice^17^. In addition to these chaperone proteins, in our recent study using co-immunoprecipitation approaches, we found that spinophilin interaction with the chaperone protein, Bip (*HSPA5*), was increased in the pancreas of obese animals^18^. However, we detected a similar number of PSMs in immunoprecipitates from high fat-diet fed wildtype and global spinophilin KO mice suggesting a non-specific interactor. The current study, utilizing UltraID and ALFA-tagged spinophilin, specifically detected Bip using both proteomics and nanobody pulldown approaches, suggesting that the use of proximity labeling and ALFA-tag nanobody pulldown may have less background and greater sensitivity than traditional co-immunoprecipitation studies.

While the pulldown of proteins involved in chaperone functions was specific when using overexpressed spinophilin, they were not selective when compared with EGFP tagged with UltraID. Moreover, as the EGFP-UltraID was expressed under control of a synapsin promoter, so the expression of this control may be less than the CMV-driven spinophilin-UltraID and therefore this interaction is probably not selective. However, even with the lack of selectivity, the expression of Hsc70 and HSC71 had a log_2_-fold change of more than –0.5 in the F451A mutant compared with wildtype spinophilin expressing cells. Therefore, at least with PP1 targeting proteins such as spinophilin, the use of non-UltraID expressing cells along with additional functional approaches, rather than an UltraID overexpressing cell is more appropriate, as the latter may lead to filtering out valid, functional interactors.

### Spinophilin interaction with proteins involved in translation and metabolism

In addition to chaperone proteins, the top 2 “Process” pathways when all specific proteins were searched were “cytoplasmic translation” and “translation.” The main proteins that were detected within this domain were elongation factors and ribosomal subunits. Previous studies determined that spinophilin modulates the function of p70 S6-kinase, a kinase that phosphorylates the ribosomal S6 subunit^41^. Herein, we found 6 elongation factor subunits that met our criteria for >4 PSMs with elongation factor 1-alpha (*EEF1A1*) as the most abundant by PSMs, averaging 51 PSMs in the two experimental conditions. This is 14.5-fold more than in the control conditions. While we detected multiple elongation factors in our EGFP-UltraID studies, we found that EEF2, EEF1G, and EEF1A1 had robust log_2_ fold-change in expression in the F451A compared with wildtype overexpressing cells (–1.21, –0.6, and –0.49 log 2-fold-changes, respectively), suggesting that wildtype spinophilin may modulate EEF expression.

In addition to translation factors, we specifically detected 61 total ribosomal subunits in our streptavidin pulldowns from spinophilin UltraID-transfected cells. The most abundant ribosomal protein was the 60S ribosomal protein L4, which had 75 PSMs across the two experimental conditions and 0 in the control samples. This recapitulates our previous studies that found a specific association of spinophilin with multiple ribosomal subunits when a goat, but not a mouse, antibody was used^17^. In that previous study, RPS3 and RPS3A had 5 and 6 PSMs, respectively, in the immunoprecipitation with the spinophilin goat antibody from wildtype mice and 0 PSMs when using the KO control^17^. However, in that study, using a mouse antibody, while there were 2 and 4 PSMs in the experimental condition matching the RPS3 and 3A, there were 4 and 3 PSMs, respectively, in the control. In the current study, there was an average of 33 and 20 PSMs matching these subunits, respectively in the spinophilin-UltraID pulldown compared to spinophilin alone expressing cells. We validated the association of spinophilin with RPS3 using a neutravidin pulldown and immunoblotting. However, the RPS3 protein in the ALFA-tag nanobody pulldown was barely detectable. This suggests that these associations may be more readily detected using proximity labeling-based approaches. In addition to translation, “Ribosome” was the top association when all spinophilin associated proteins were probed using the Kyoto Genes and Genomes pathways analysis (KEGG), again demonstrating a critical association of spinophilin with ribosomal proteins. While specific, again we detected ribosomal proteins in the EGFP-UltraID pulldowns. However, even after subtracting the non-selective interactors, we still found that translation was one of the top biological processes. Moreover, we detected 11 ribosomal subunits that were decreased in the F451A spinophilin mutant overexpression compared to wildtype overexpressing cells, suggesting a functional regulation of these proteins by spinophilin.

### Spinophilin limits its own expression but stabilizes proteins associated with ribosome, RNA binding, and translation

Previous studies using microdissections of neuropil and cell-body enriched areas of the hippocampus, suggest that spinophilin is one of the most highly translated mRNAs within synaptic regions^30,31^. Using RNAScope, we found that spinophilin mRNA is highly neuropil localized in the striatum, when compared with the spinophilin homolog, neurabin. Given our findings that spinophilin interactions with, and possible promotion of, the expression of RNA binding proteins and proteins involved in mRNA translation, we posit that spinophilin’s neuropil RNA localization may underlie its unique regulation of striatal functions when compared with its homolog neurabin^12^.

We previously generated ROSA-targeted, Cre-dependent, HA epitope-tagged spinophilin knock-in mice. Unfortunately, HA-spinophilin was lowly expressed, ∼1.5% of endogenous, using this approach. While we attributed this low expression to using the endogenous ROSA promoter^16^, herein, we found that injection of a virus encoding a Cre-dependent, HA-spinophilin under control of a strong CMV promoter also resulted in minimal spinophilin overexpression. However, when we injected conditional spinophilin KO mice with this construct, we observed robust spinophilin expression when spinophilin was previously knocked out of those cells. While spinophilin limits its own expression, previous studies have found that loss of spinophilin decreases PP1 expression^20,42^. Taken together, these data suggest that *in vivo* overexpression studies involving spinophilin are severely limited and viral-mediated overexpression approaches will most likely not recapitulate the neuropil-enriched localization of spinophilin mRNA. To overcome these shortcomings, future studies using transgenic mice that Cre-dependently express an endogenous ALFA-UltraID tagged spinophilin or emerging CRISPR-based viral approaches to endogenously tag the spinophilin locus with a proximity label and epitope-tagged sequences will need to be leveraged^43^.

Importantly, we found that loss of spinophilin decreased the PSD expression of known spinophilin interacting proteins^15,16,20,39^, including PSD-95, GluN2B, GluA2, and two PP1 isoforms. Interestingly, *Dlg4* (PSD95) mRNA is enriched in the hippocampal neuropil compared to somatic compartment^30^. Conversely, *Gria2* (GluA2) mRNA is enriched in the hippocampal somatic compartment compared to the neuropil^30^. The local translation of *Grin2b* (GluN2B) is unclear as it was not listed in the paper by Schuman and colleagues; however, Grin2d was found to be neuropil enriched^30^. To this end, GluA2 was not PSD enriched in our fractionation whereas PSD95 and GluN2B were. Moreover, in the synaptoneurosome fraction of spinophilin KO compared with wildtype mice, GluA2 did not have a decreased expression whereas both PSD95 and GluN2B trended towards a decreased expression. Interestingly, we observed either a strong trend or significant decreases in total protein expression within the synaptoneurosome and PSD fractions when normalized to the total protein in the homogenate. Therefore, loss of spinophilin globally decreases protein expression within these fractions. Importantly, most of the proteins had no change in expression within the synaptoneurosome or PSD fractions when normalized to total protein abundance. The exceptions to this were the PP1 isoforms. We found that there was a trend or significant decrease in the two PP1 isoforms within the synaptoneurosome and/or PSD fractions even when normalized to the decreased PSD protein expression. This may suggest that loss of spinophilin limits PP1 targeting to the PSD. Overall, these data suggest that loss of spinophilin may mediate local neuropil mRNA translation and that loss of spinophilin is globally decreasing PSD protein expression or modulating the organization of the PSD such that it is not biochemically fractionating in the same way as in wildtype mice.

### Spinophilin interactions and disorders associated with mRNA translation

Many neurodevelopmental disorders are associated with excessive neuropil mRNA translation. For instance, loss of the FMRP protein, that is the cause of the neurodevelopmental Fragile X syndrome, leads to excessive mRNA translation of specific target genes^44^. In our HEK293 cell studies, we detected 13 out of 121 proteins within the fragile X pathway using WikiPathways analysis. These include the major Fragile X-associated protein, synaptic functional regulator Fmr1 (Fmrp; *FMR1*). We detected an average of 10 PSMs across the two samples and 0 PSMs across all the spinophilin and EGFP-UltraID control samples. Fmrp is a protein that can act as a translational repressor and can impact mRNA splicing^45^. The *PPP1R9B* gene is genetically linked to neurodevelopmental disorders^46–48^. Moreover, our recent studies demonstrate that spinophilin limits grooming in a genetic mouse model for excessive grooming^10^. Therefore, future studies will need to probe how spinophilin mediates protein synthesis changes within the context of neurodevelopmental disorders.

## Supporting information

Supplemental Westerns

Scaffold Files

Genbank Files

## Acknowledgements

The authors acknowledge grant support from NIH – R21AA030319 (Baucum and Atwood), NIH-R33DA041876 (Baucum), Cagiantas Fellowship (Hens), Startup support from Department of Pharmacology and Toxicology, Stark Neurosciences Research Institute, and Strategic Recruitment Initiative, Indiana University School of Medicine (Baucum).

The mass spectrometry work performed in this work was done by the Indiana University School of Medicine Center for Proteome Analysis. Acquisition of the IUSM Center for Proteome Analysis instrumentation used for this project was provided by the Indiana University Precision Health Initiative. The proteomics work was supported, in part, by the Indiana Clinical and Translational Sciences Institute (UL1TR002529 from the National Institutes of Health, National Center for Advancing Translational Sciences, Clinical and Translational Sciences Award) and the P30 IU Simon Comprehensive Cancer Center Support Grant (P30CA082709).

## Materials and Methods

### Buffer, reagent, and equipment information

Complete information about buffer components, reagents, antibodies, equipment, and software including names and catalog numbers and company information are given in **Tables SM1-SM5**.

### Generation of ALFA-UltraID Constructs

The vector pAAV.CMV.PI.EGFP.WPRE.bgh^49^ was digested with NotI and BamHI to excise the EGFP sequence. We next generated 3 DNA fragments with overlap to the vector backbone or with each other (**Table SM6)**. We assembled the fragments and vector using NEBuilder HiFi DNA Assembly Master Mix using manufacturer’s instructions. Miniprep DNA was screened by PCR and positive DNAs were sent for whole plasmid sequencing using Oxford Nanopore Technology with custom analysis and annotation (Plasmidsaurus, Eugene, OR). The Genbank file for a positive sequence is given in supplemental file “SpinoLoxp8.gbk”. It is important to note that UltraID is defined as amino acids 2-171 of the full-length sequence in the construct and that there is an additional 30 amino acids at the C-terminus of the sequence that were included in our constructs as they were part of the plasmid sequence.

We also created an EGFP with a C-terminal UltraID protein. This construct was generated in a construct using a human synapsin promoter (pAAV-hSyn-EGFP) to permit viral expression in mammalian neurons; however, it also expresses in HEK293 cells^27^. The Genbank file for a positive sequence is given in supplemental file “Claeboe_Y3S_1_GFP-UID_pLann.gbk”.

### Transfection of HEK293 cells

HEK293FT cells were used for mammalian protein expression. Originally cells were purchased and split into passage 8 and 9, respectively, and frozen down for long term storage. Thawed cells were allowed to recover for 24-48 hrs in Recovery Media prior to beginning switched into Complete Media Cells were allowed 1 week of growth in tissue culture incubator at 37°C and 5% CO_2_ until next passage into maintenance T-75 flask and experimental 6-well plates. During passage cells were counted and density adjusted to 300,000/mL in 2mL for 6-well plates per Falcon seeding suggestion. Transfections of cells were conducted 48 hours post cell passage wherein cell displayed 60-70% confluency. Cells were transfected using Lipofectamine 3000 using 5 μg of DNA (Spinophilin Loxp, D2-eGFP, and Cre/pDonr vector) using the manufacturer’s protocol.

### Biotin Treatment

Forty-eight hours post transfection, cell medium was aspirated off and cells were washed 3x with Buffer B. Cells were incubated in 50 μM Biotin diluted in Buffer B for 3.5 to 4 hours at 37°C. Cells were washed 2 times with Buffer B to remove any free, unreacted biotin.

### HEK293 cell lysis and affinity purification

Cells were lysed by homogenization in 750 µl RIPA Buffer. Homogenized lysates were then sonicated (20% amplitude for 15 seconds). Cell lysates were then rotated for 30 min at 4°C. Cellular debris was pelleted by centrifugation (14,000 x g, 15 min, 4°C). 100 µl of sample was mixed with 33.3 µl 4X sample buffer for an input and 600 µl was used for affinity purification using 20-40 µl of Neutravidin, trypsin-resistant streptavidin, or ALFA-selector beads (20 – 40 µl) and allowed to rotate overnight at 4°C. The following day, beads were washed 1x in RIPA Buffer followed by 2x in IP wash buffer or 3 x in IP wash buffer with 5 min of rotation at 4°C between each wash step. Following washes, samples were eluted in 2X SDS sample buffer for Western Analysis or washed 2 additional times in PBS for mass spectrometry. For PBS washes, samples were centrifuged rather than separated by the magnet as they did not stick as well to the magnet when switching buffers to PBS.

### Immunoblotting

Cell lysate inputs and/or pulldown samples were heated at 70°C for 10 min and vortexed both before and after heat. The beads within the pulldown samples were pulled from solution through either a magnetic rack or centrifugation (2,000xg for 1 min) for the Streptavidin and Neutravidin beads, respectively. Fifteen μL of either input or pulldown sample was loaded onto a precast 10% Criterion Gel. Gel electrophoresis was performed and total protein staining with Revert stain, and immunoblotting was performed as we have previously described^9,10,15,16,20^. Immunoblots were imaged using an Odyssey M machine and acquired with Odyssey acquisition software. Images were linearly adjusted for brightness and contrast and exported using Image Studio software (LI-COR Biosciences). When quantifications were performed, the band intensity from input samples were normalized to a Revert 520 total protein stain.

### Proteomics studies: Streptavidin on bead digests

Sample preparation, mass spectrometry analysis, bioinformatics, and data evaluation for quantitative proteomics and phosphoproteomics experiments were performed in collaboration with the Indiana University School of Medicine Center for Proteome Analysis similarly to several previously published protocols.^50,51^. After washing, beads were covered with 8 M Urea, 100 mM Tris hydrochloride, pH 8.5, reduced with 5mM tris (2-carboxyethyl) phosphine hydrochloride (TCEP) for 30 minutes at room temperature to reduce the disulfide bonds. The resulting free cysteine thiols were alkylated using 10 mM choloracetamide (CAA) for 30 minutes at RT, protected from light. Samples were diluted to 2 M Urea with 50 mM Tris pH 8.5 and proteolytic digestion was carried out with Trypsin/LysC Gold (0.4 µg) overnight at 35 °C. After digestion, samples were quenched with 0.4% trifluoroacetic acid (v/v).

### Proteomics studies: Streptavidin pulldown LC-MS

Approximately 1/15th of each IP sample was loaded onto a 5 cm C18 trap column Acclaim™ PepMap™ 100 (3 µm particle size, 75 µm diameter) followed by a 25 cm EASY-Spray column and analyzed using a Q-Exactive Plus mass spectrometer operated in positive ion mode. Solvent B was increased from 5%-35% over 100 min, to 90% over 2 min, back to 3% over 2 minutes (Solvent A: 95% water, 5% acetonitrile, 0.1% formic acid; Solvent B: 100% acetonitrile, 0.1% formic acid). A data dependent top 15 method was used with MS scan range of 350-2000 m/z, resolution of 70,000, AGC target 3e6, maximum IT of 200 ms. MS2 resolution of 17,500, scan range of 200-2000 m/z, normalized collision energy of 30, isolation window of 4 m/z, target AGC of 1e5, and maximum IT of 150 ms. Dynamic exclusion of 10 sec, charge exclusion of 1, 7, 8, >8 and isotopic exclusion parameters were used.

### Proteomics studies: Streptavidin pulldown data analysis

Data were analyzed using Proteome Discoverer 2.5.0.400 (Thermo Fisher Scientific). A *Homo sapiens* reference proteome database (UniProtKB/TrEMBL; 78806 sequences downloaded 05132022), plus UltraID tagged spinophilin sequences plus common laboratory contaminants (73 sequences including streptavidin) was searched using SEQUEST HT. Precursor mass tolerance was set to 10 ppm and fragment mass tolerance set at 0.02 Da with a maximum of 3 missed cleavages. A maximum of 3 modifications were allowed per peptide Dynamic modifications include methionine oxidation; biotin on lysine, phosphorylation on serine, threonine, and tyrosine; dynamic protein terminus modifications were acetylation, met-loss, and met-loss plus acetylation. Static modifications were carbamidomethylation on cysteines. Percolator false discovery rate (FDR) filtration of 1% was applied to both the peptide-spectrum match and protein levels. Search results were loaded into Scaffold Q + S Software (version 5.2.2, Proteome Software, Inc) for visualization. Scaffold files for spinophilin interactome (2024_03_78_spino.sf3 and 2024_03_78_spino_2.sf3, without and with searches for spinophilin UltraID and streptavidin, respectively) were uploaded as a supplementary file. Scaffold files for EGFP UltraID (2024_09_247_eGFP.sf3) was uploaded as a supplementary file.

### Proteomics studies: TMT proteomics experiment, labeling

Cell pellets were lysed in 200 µL of 8 M Urea, 100mM Tris hydrochloride, pH 8.5 by 30 rounds of high power 30sec on 30 sec off in a Diagenode Bioruptor sonicator. Samples were clarified by centrifugation at 12,000 rcf for 20 min and protein concentration determined by Bradford Assay (Biorad). Approximately 30 µg of protein from each sample was then reduced with 5mM TCEP for 30 minutes at room temperature and alkylated using 10 CAA for 30 minutes at RT, protected from light. Samples were diluted to 2 M Urea with 50 mM Tris pH 8.5 and proteolytic digestion was carried out with Trypsin/LysC Gold (1:25 protease to substrate) overnight at 35 °C. After digestion, samples were quenched with 0.4% TFA. Peptides were cleaned up using Waters Sep-Pak® Vac cartridges with a wash of 1 mL 0.1% TFA followed by elution in 0.6 mL of 70% acetonitrile 0.1% formic acid (FA). Peptides were dried by speed vacuum and resuspended 50 mM triethylammonium bicarbonate. Each sample was then labeled for 2 hours at room temperature with 0.5 mg Tandem Mass Tag Pro (TMTpro) reagent; label tables for both included below. Samples were checked to ensure labeling efficiency of >90 % and then quenched with 0.3% hydroxylamine (final v/v) at room temperature for 15 min. Labeled peptides were then mixed and dried by speed vacuum.

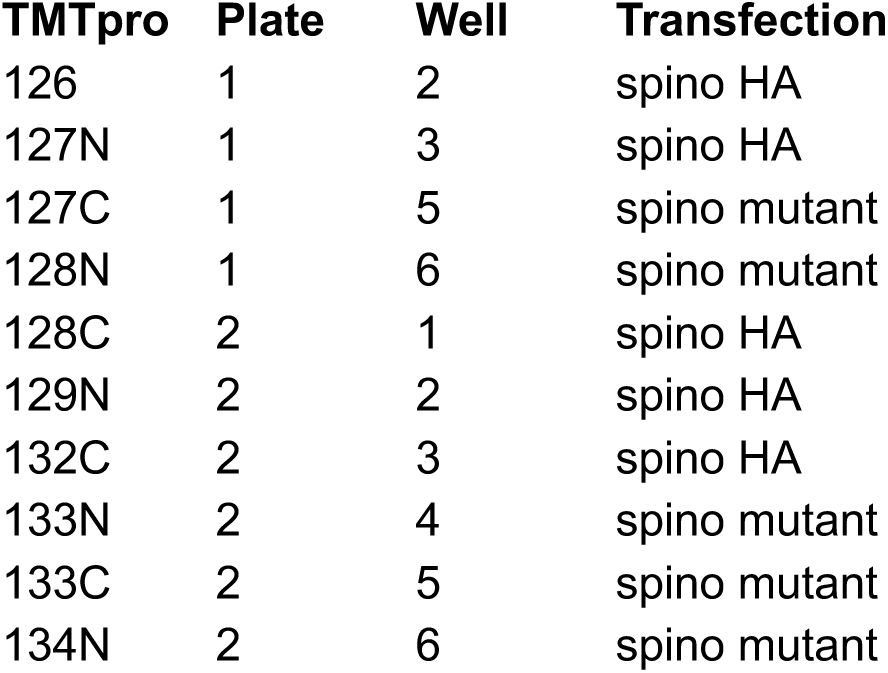

### Proteomics studies: TMT proteomics experiment, High pH Basic Fractionation

Approximately 1/3^rd^ of the labeled peptide mixture was fractionated using the TMT fractionation protocol of Pierce high pH basic reversed-phase peptide fractionation kit; with a wash of 5% acetonitrile, 0.1% triethylamine (TEA) followed by elution in 10%, 12.5%, 15%, 17.5%, 20%, 22.5%, 25%, 30%, and 70% acetonitrile, all with 0.1% TEA.

### Proteomics studies: TMT proteomics experiment, LC-MS/MS

1/5^th^ of each global peptide fraction was injected using an EasyNano LC1200 HPLC coupled with 25 cm Aurora column on an Eclipse Orbitrap mass spectrometer with FAIMSpro installed. Peptides were eluted over a 180-minute method: Solvent B was increased from 5%-30% over 160 min, to 85% B over 10 minutes, and down to 10% B. The mass spectrometer was operated in positive ion mode, advanced peak determination on and Easy-IC™ used, with 3 FAIMS CVs (–50, –60, –70). A cycle time of 2 s was used for each CV and RF lens was set to 30%. MS1 parameters for each cycle were: orbitrap resolution of 120,000, scan range of 400-1600 m/z, standard AGC, 50 ms max IT, minimum intensity of 2.5e4, charge state 2-6, 60 sec dynamic exclusion. MS2 settings were quadrupole isolation of 0.7 m/z, fixed HCD of 34, orbitrap resolution of 50,000, fixed first mass of 100, 200% AGC, dynamic max IT.

### Proteomics studies: TMT proteomics experiment, LC-MS/MS data analysis

Data were analyzed in Proteome Discoverer 2.5 using a *Homo sapiens* reference proteome UniProt FASTA (downloaded 10/04/2019) sequences plus common contaminants (71 sequences) as well as HA or UltraID spinophilin, UltraID-EGFP, streptavidin, or GFP-DRD2 protein sequences. SEQUEST HT searches were conducted with a maximum number of 3 missed cleavages; precursor mass tolerance of 10 ppm; and a fragment mass tolerance of 0.02 Da. Static modifications used for the search were, 1) carbamidomethylation on cysteine (C) residues; 2) TMT label on lysine (K) residues 3) TMT label on the N-termini of peptides. Dynamic modifications used for the search were oxidation of methionines, phosphorylation on serine, threonine or tyrosine, deamidation on arginine, and acetylation, methionine loss or acetylation with methionine loss on protein N-termini. Percolator False Discovery Rate was set to a strict setting of 0.01 and a relaxed setting of 0.05. IMP-ptm-RS node was used for all modification site localization scores. Values from both unique and razor peptides were used for quantification. In the consensus workflows, peptides were normalized by total peptide amount with no scaling. Quantification methods utilized isotopic impurity levels available from Thermo Fisher Scientific. Reporter ion quantification was allowed with S/N threshold of 6 and co-isolation threshold of 30%. Resulting grouped abundance values for each sample type, abundance ratio (AR) values; and respective p-values (individual protein, ANOVA) from Proteome Discover™ were exported to Microsoft Excel for downstream pathway analysis using String-db.

### Proteomics data availability

Raw and processed mass spectrometry data have been uploaded to the ProteomeXchange partner MassIVE with repository ID MSV000096575. Data will be accessible upon publication.

### RNAScope

Eight-week-old Male and Female Neurabin Flox/+ mice were perfused transcardially with 10 mL PBS, followed by 4% paraformaldehyde in PBS at a rate of 1 ml/min. Brains were removed and stored overnight in 4% paraformaldehyde for 24 hours at 4⁰C. Brains were then immersed 10%, 20%, and 30% sucrose in 1x PBS at 4⁰C for 24 hours each before being frozen with dry ice and embedded in TFM media. Striatal cryostat sections (10 µm) were mounted on Colorfrost Plus slides and taken through the RNAScope protocol for fixed frozen tissue samples, analogous to formalin fixed paraffin embedded (FFPE) as previously described [13]. Tissue sections were washed in 1x PBS for 5 minutes, post fixed for 15 min at 4⁰C in 4% PFA in PBS. Following an ethanol series for dehydration (50% ethanol for 5 minutes, 70% ethanol for 5 minutes, 100% ethanol for 5 minutes, 100% ethanol for 5 minutes), tissue sections were treated with RNAscope Hydrogen Peroxide for 10 minutes at room temperature (RT) and incubated in target retrieval reagent for 15 minutes maintained at a temperature of 100-103⁰C and immediately rinsed in deionized water. Following placement of a hydrophobic barrier around the tissue section with a hydrophobic pen, sections were treated with Protease III for 30 minutes at 40 ⁰C in the HybEZ hybridization oven and washed with de-ionized water twice at RT, for two minutes each time, prior to proceeding to probe hybridization. For probe hybridization, the tissues were then incubated with target probes for Spinophilin or Neurabin in probe diluent for 2 hours at 40⁰C; slides were then incubated with AMP1 for 30 minutes at 40⁰C, AMP2 for 30 minutes at 40⁰C, and then AMP3 for 15 minutes at 40⁰C. AMP1-3 were from the RNAscope Multiplex Fluorescent Detection Reagents V2. After each step, slides were washed twice at RT, for two minutes each, with RNAScope Wash Buffer. For multiplex detection, tissues were incubated with 4-6 drops of horseradish peroxidase of the probe corresponding channel for 15 minutes at 40⁰C; washed with RNAScope Wash Buffer 2 minutes at RT twice; and incubated with 150 microliters of Opal 690 or Opal 520 fluorophores diluted in TSA buffer for 30 minutes at 40⁰C. Subsequently, these tissues were washed twice with RNAScope Wash buffer 2 minutes at RT twice; and incubated with HRP blocker at 40⁰C; and washed with RNAScope Wash Buffer for 2 minutes at RT twice. This process was repeated for each probe. To counterstain, slides incubated with 4 drops of DAPI for 30 seconds at RT and immediately incubated with ProLong Gold Anitfade Mounting Mountant. Coverslips were placed on tissue sections, sealed with nail polish and allowed to dry overnight prior to imaging.

### Imaging

To generate the results in Figure 6A, widefield fluorescence imaging for spinophilin and neurabin was performed on the Cytation5 plate reader/microscope using a 4X objective, and composite images were generated using the image stitching function. Spinophilin mRNA or neurabin mRNA probe was detected with Opal690 fluorophore in separate sections.

To generate the results in Figure 6B, microscopy was performed on a Zeiss LSM 900 with Airyscan 2. Spino mRNA was detected with Opal520 fluorophore while or neurabin mRNA was detected with Opal690 fluorophore within the same section. Dorsal and ventral striatal subregions were chosen in correspondence with the Paxinos and Franklin Mouse Brain Atlas and imaged using the Plan-Apochromat 63x/1.40 Oil objective, obtaining optical sections (pixel size = 0.05 micron x 0.05 micron). Optical sections were captured using Airyscan functionality. Individual channels were imported into Photoshop (Adobe, San Jose, CA) and levels linearly adjusted for picture presentation. Pictures were assembled in Illustrator (Adobe).

### Virus generation

HA epitope-tagged spinophilin was cloned into pAAV-CBh-DIO-EGFP using AscI forward primer (CGCGCCTTATCTAGATTTAGCCACC ATGTACCCATACGACGTCCCAGACTATGCT ATGATGAAGACGGAGCCAC) and NheI reverse primer (CTAGCGCTACCGGTCGCCACCGCCACCATGTACCCATACGACGTCCCAGACTATGCT ATGATGAAGACGGAGCCAC). Cre-dependent, HA-tagged spinophilin expression was validated using HEK293 cells and viral constructs were sent to Vigene Biosciences (Rockville, MD, now a part of Charles River) for virus generation. AAV2/9 virus was generated by Vigene and shipped at a titer of 1.24×10^13GC/mL.

### Virus transduction

Mice were anesthetized with 5% isoflurane for 5 minutes (flow rate = 2 L/min). The hair on top of the mouse’s skull was trimmed and the mouse nose was secured into nose cone. Once secured, sterile ocular lubricant was added to mouse eyes and the isoflurane was lowered to 1.5%. Ear bars were inserted into mouse ear canals to stabilize skull, then skull was sterilized with ethanol and iodine successively at least three times each. Once skull was secured and leveled, a scalpel was used to make incision down the base of the skull long enough to expose bregma and lambda markings on skull. The skull was next leveled along the X– and Y-axes using Alignment Indicator, then a microscope was used to locate the center of bregma, then all stereotax coordinates on a Micro Manipulator with Digital Display were re-zeroed. A stereotax drill was used to create two bilateral holes (4 total) in the skull to infuse anterior and posterior regions of the striatum with AAV. Specifically, drill holes for anterior striatal infusions were placed at: anterior/posterior (A/P) +1.00, 5medial/lateral (M/L) +/-2.05, dorsal/ventral (D/V) –3.5, and drill holes for posterior striatal infusions were placed at: A/P +0.4, M/L +/-2.65, D/V –3.55. Syringe was attached to stereotax and enough AAV for each injection (one at a time) was withdrawn into syringe, such that 150nL was injected into anterior drill holes and 300nL was infused into posterior drill holes. Once syringe was loaded with AAV, the syringe was aligned with appropriate drill hole and was quickly lowered to appropriate Z-coordinate where AAV was infused at a rate of 40 nL/min. After infusion, the AAV diffused undisturbed for 5 minutes, then the syringe needle was slowly removed from skull. Between injections, the syringe needle was wiped with isopropanol pad and rinsed by pipetting ethanol and milli Q water onto KimWipes. After all injections were completed, the skin on skull was super glued back together with tissue adhesive and the mouse received a 5 mg/kg ketoprofen and 0.5 ml of warm saline via a subcutaneous injection. Finally, anti-itch cream was applied to skull and mouse was removed from ear bars and placed into pre-warmed biohazard cage. Animals were weighed and treated with anti-itch cream inside a biological safety cabinet once a day for 72 hours following surgery, then animals could return to standard housing.

### Subcellular fractionation

Subcellular fractionation protocol was adapted from Villasana et al^32^. All steps from dissection to isolated PSD resuspension carried out on ice with cold buffers and solutions. Four total striata were dissected from male or female spinophilin wildtype or global knockout mice were homogenized in cold synpatoneurosome buffer via mechanical glass tissue grinder. An n of 4 was used per group and striata from 2 animals per “n” value were pooled together. A homogenate fraction was set aside for western analysis prior to dilution of homogenate by 1x volume of synpatoneurosome buffer. (e.g. if volume is 8 mL add 8 mL). Further cell lysis was conducted through three rounds of sonication pulses for 20 seconds at 20% Amp 1 power. Sample was then loaded into 50 mL Luer-lok syringe for 2x filtration through pre-wetted 100 μm pore nylon filter stamped out that morning for the 13 mm diameter filter holder. Course filtered homogenate was then loaded into a 5 mL Luer-lok syringe to pass through a 5 μm pore hydrophilic filter held in the filter holder in increments of 4 mL. Due to pressure build ups it is recommended the experimenter swap the filter paper every 5-6 mL of sample filtrate. Samples pooled into a clean 50 mL polycarbonate tube prior to centrifugation at 1,000 x g for 10 min. Resulting isolated synpatoneurosome pellet (pre– and post-synapse) was resuspended in 1.25 mL resuspension buffer. A portion of the SN fraction was set aside for western analysis prior to 1x sample volume addition of Dilution Buffer. Samples allowed to stir in open top container for 15 min at 170 rpm in water ice bath prior to centrifugation at 16000 rpm for 20 min at 4°C in a benchtop ultra centrifuge. Resulting pellet was resuspended in 50 μL of resuspension buffer then layered onto a sucrose gradient created in a centrifuge tube containing equal volumes of layered Phase 1 and Phase 2 buffers. Samples were then centrifuged at 40000 rpm for 2 hrs in swinging bucket rotor at 4°C. Pellet ring was carefully collected and resuspended in 70 μL of resuspension buffer with further dilution by 70 μL of PSD dilution buffer. Samples were centrifuged for a final run at 40000 rpm for 30 min at 4°C to yield isolated PSD pellet that was then resuspended in 50 μL of resuspension buffer and stored for western analysis.

**Table SM1.**
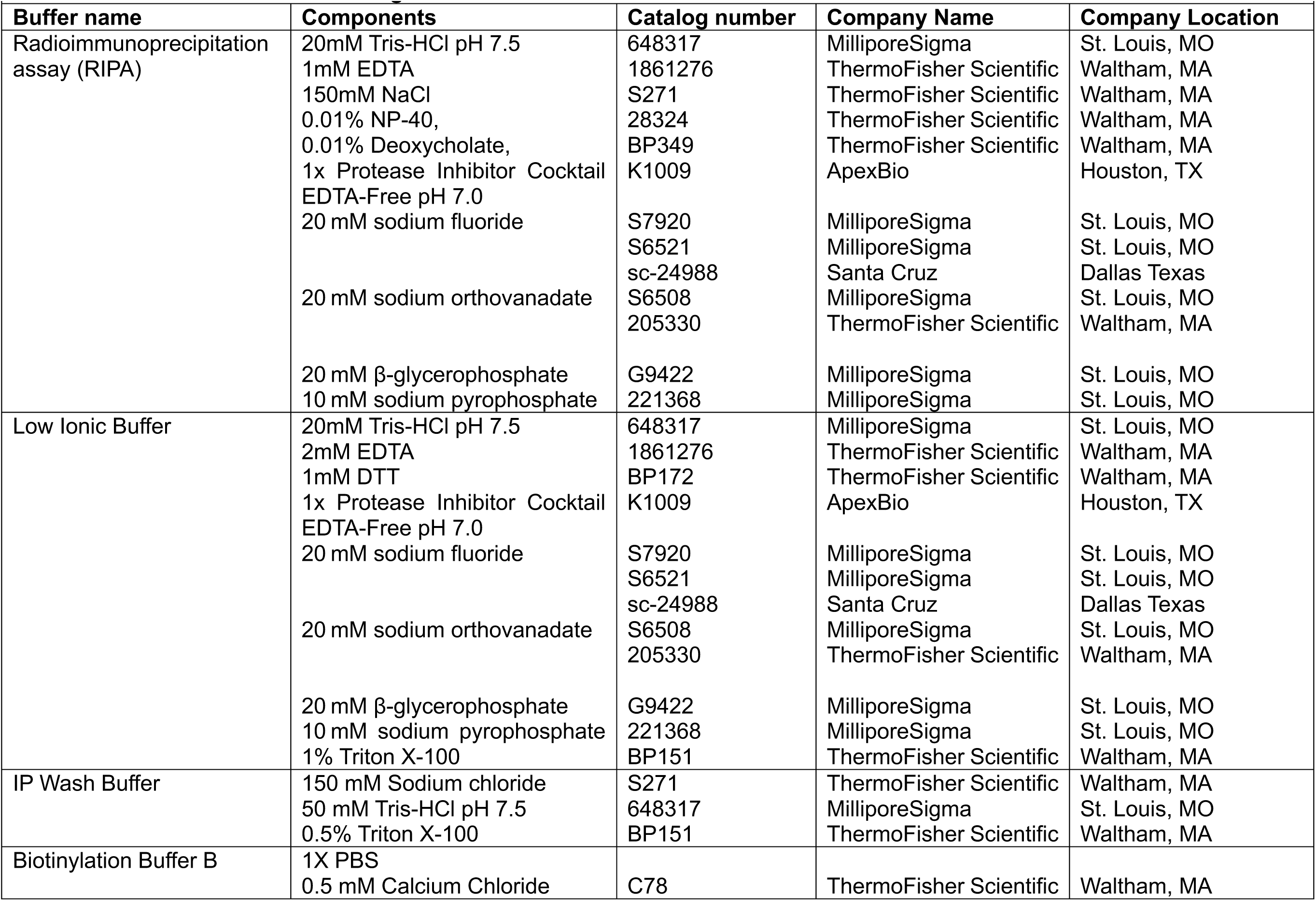
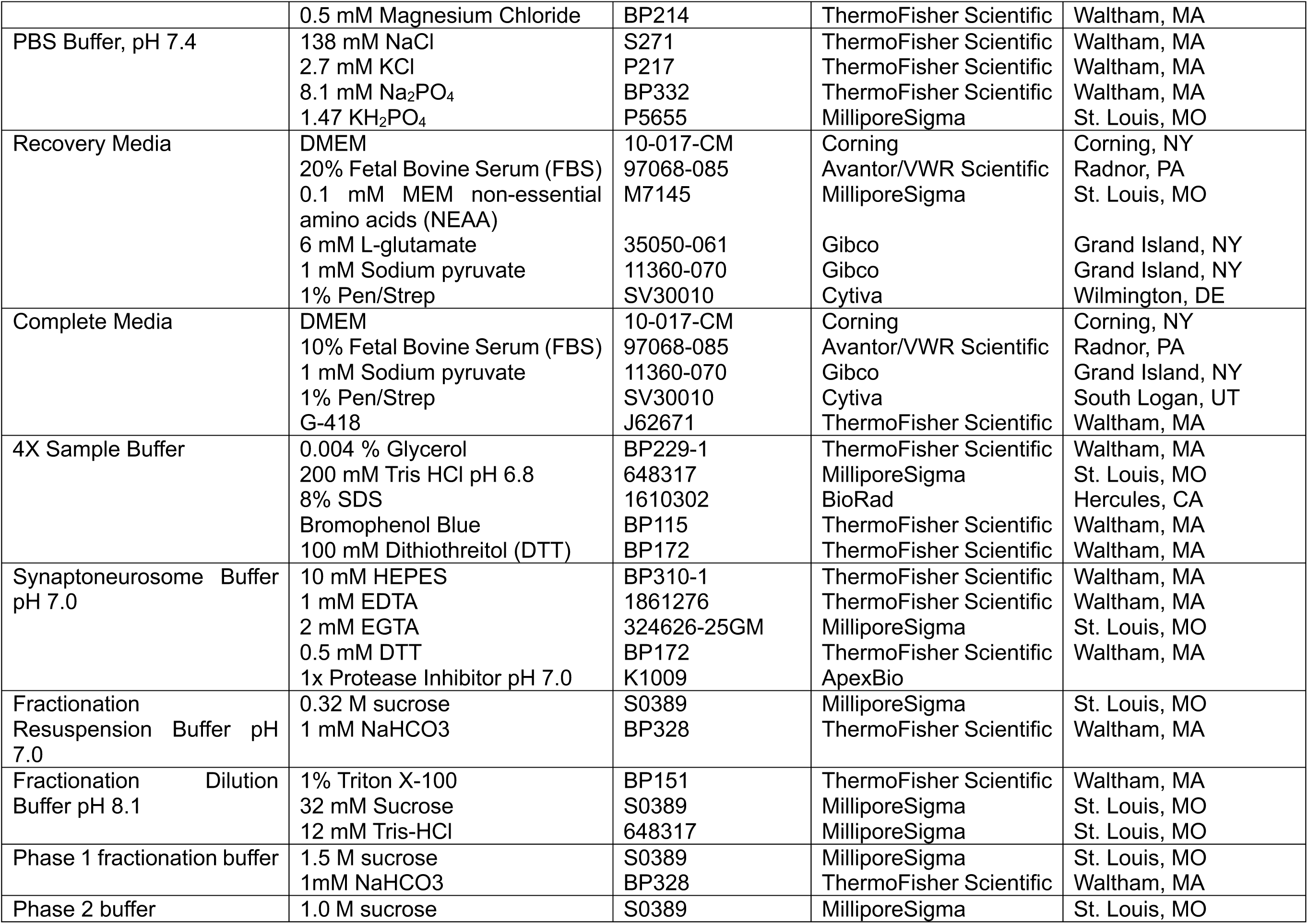
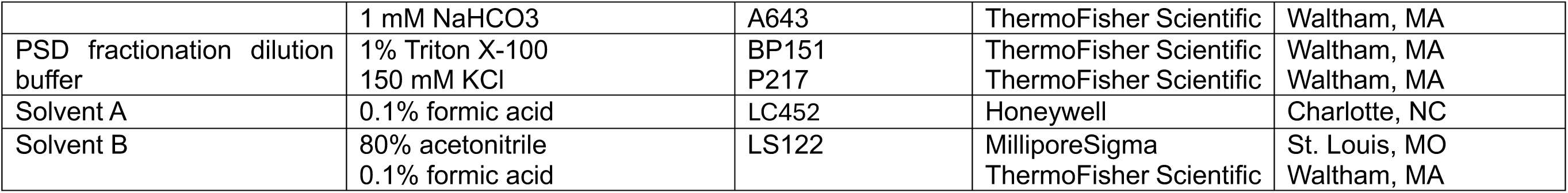
Buffers and chemical reagents.

**Table SM2.**
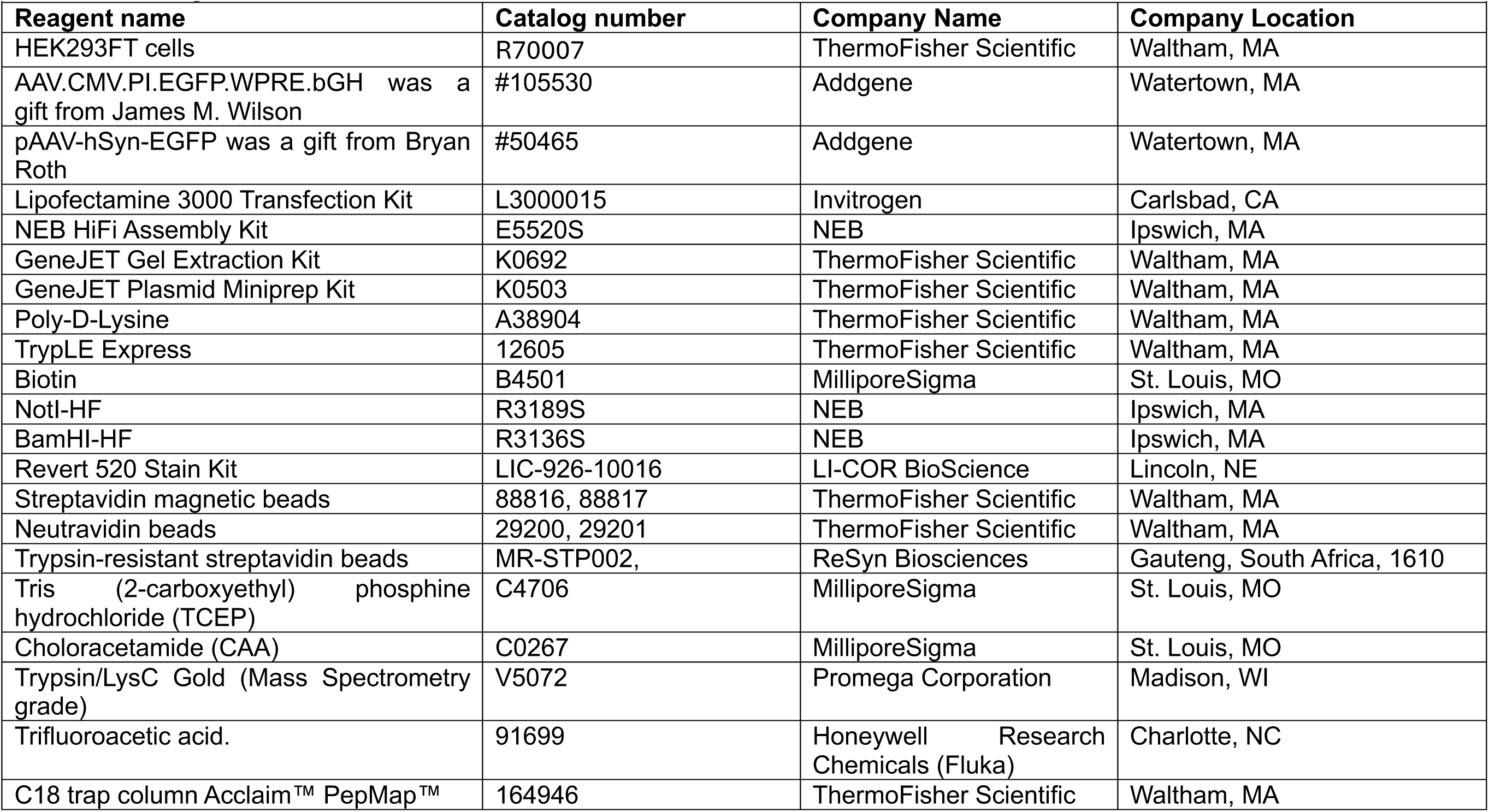
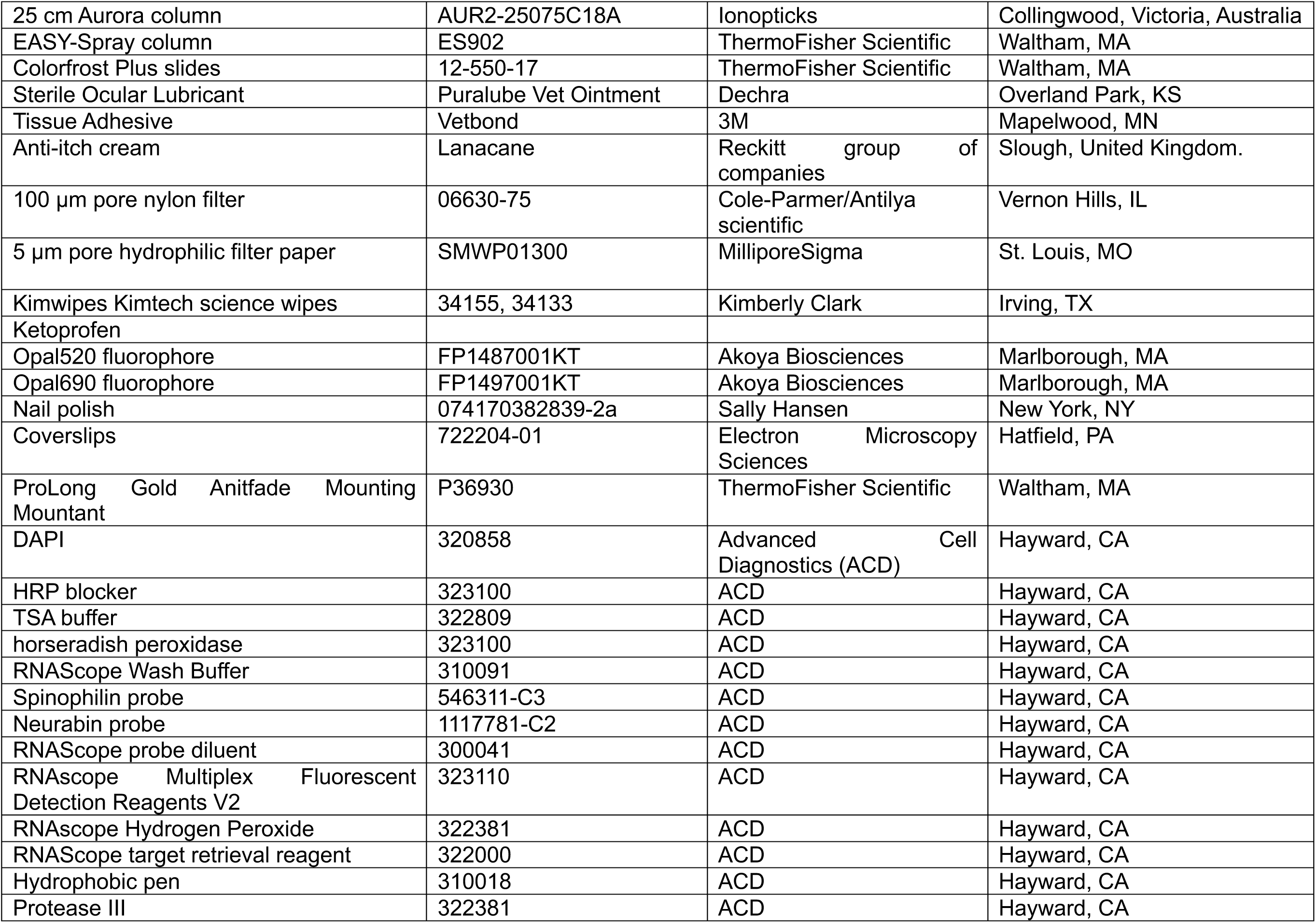
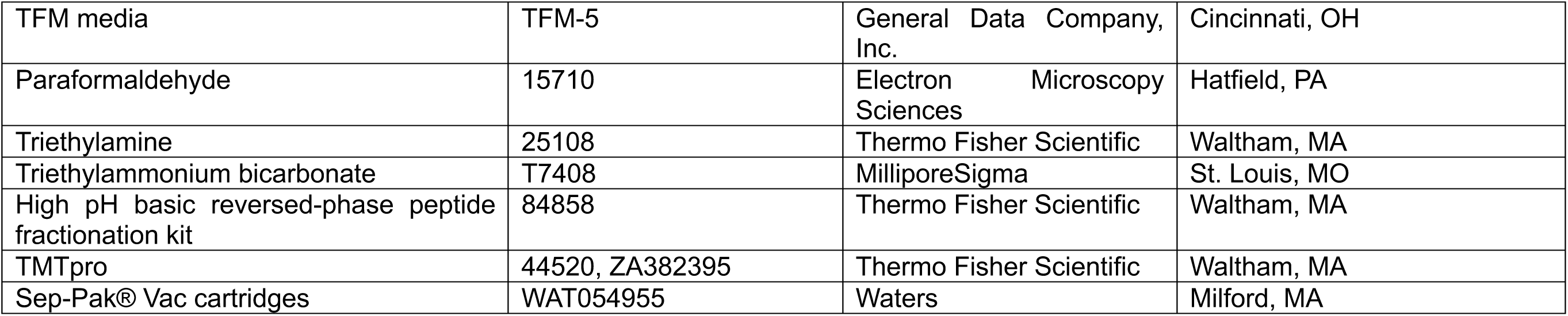
Reagents.

**Table SM3.**
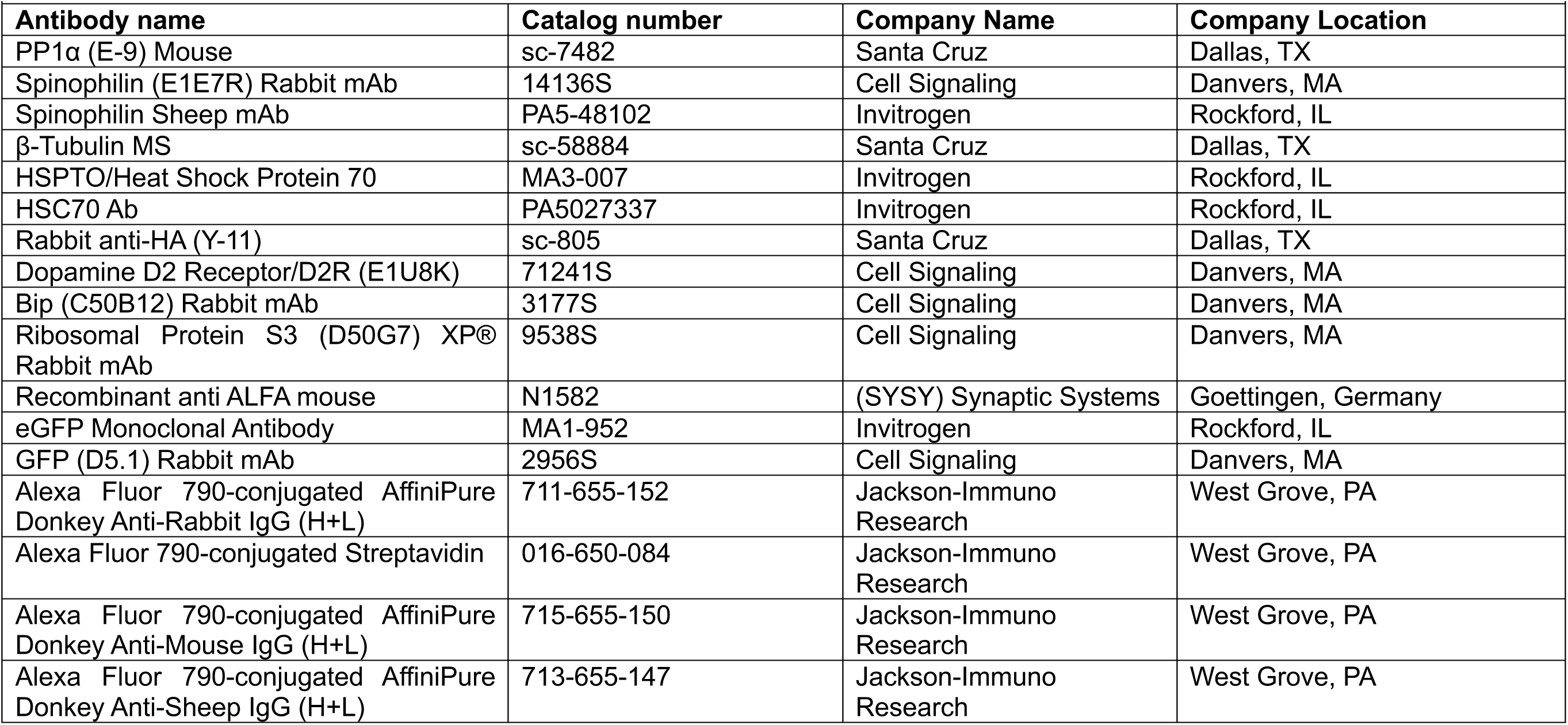
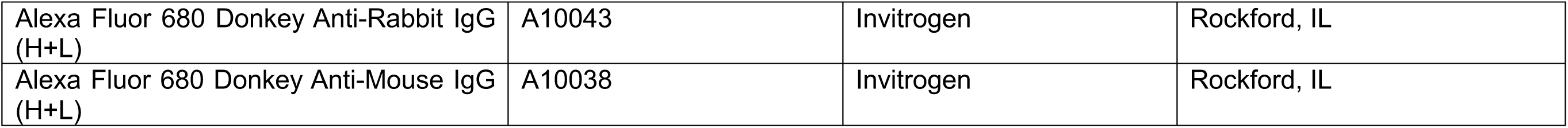
Antibodies.

**Table SM4.**
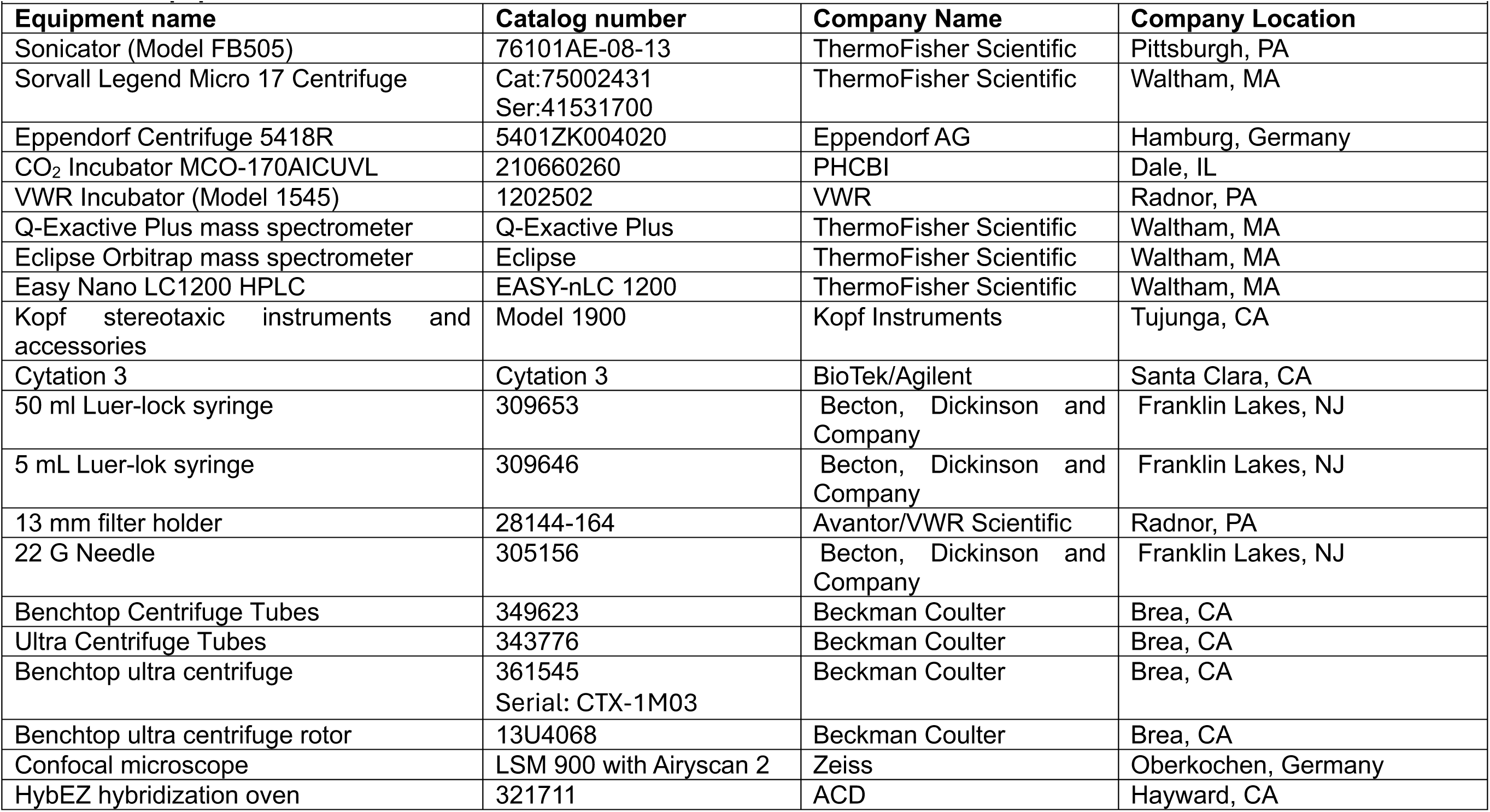
Equipment.

**Table SM5.**
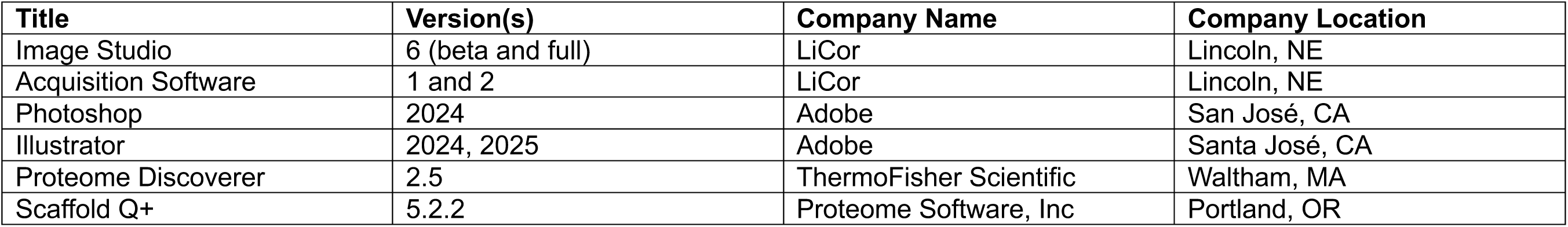
Software.

**Table SM6.**
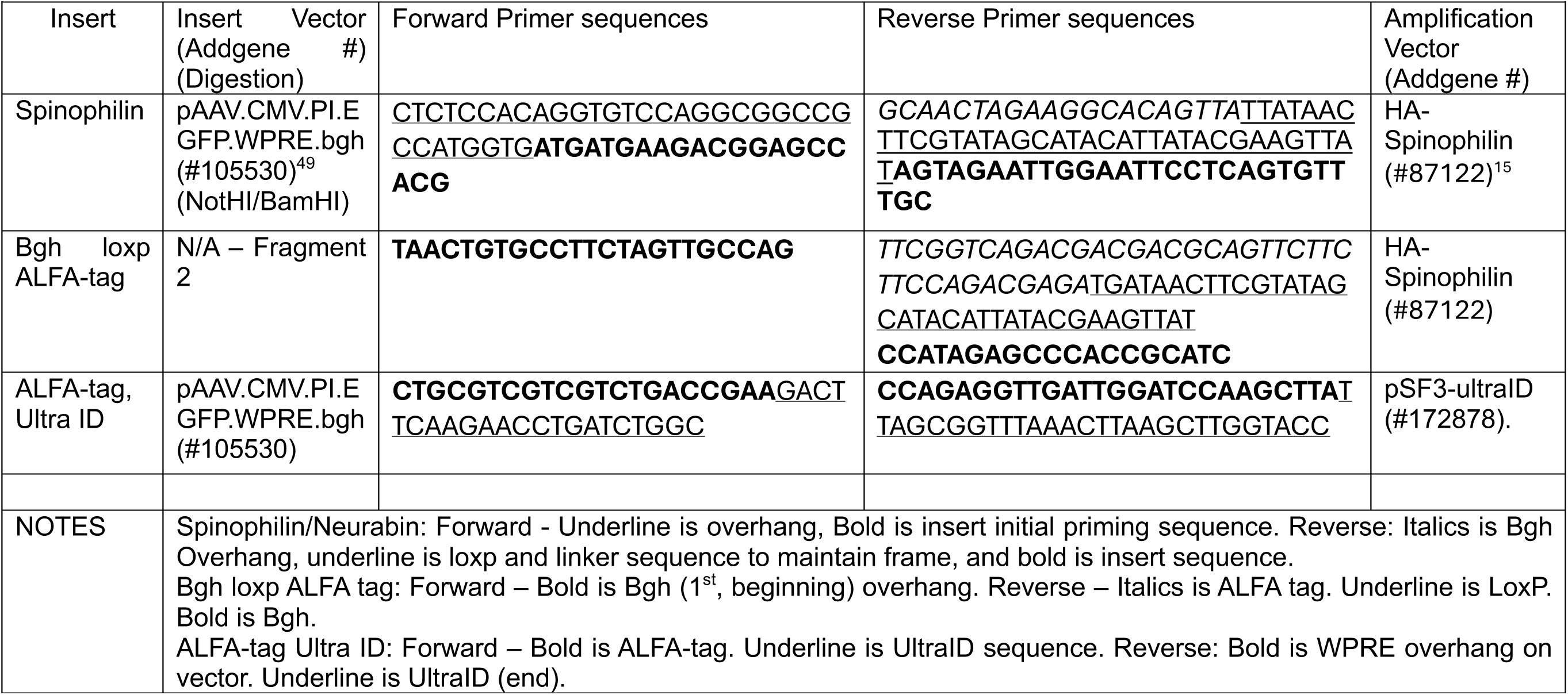
Primer sequences.

