## Supplementary figures and images for "Proximity labeling and orthogonal nanobody pulldown (ID-oPD) approaches to map the spinophilin interactome uncover a putative role for spinophilin in protein homeostasis"

### Figure S1

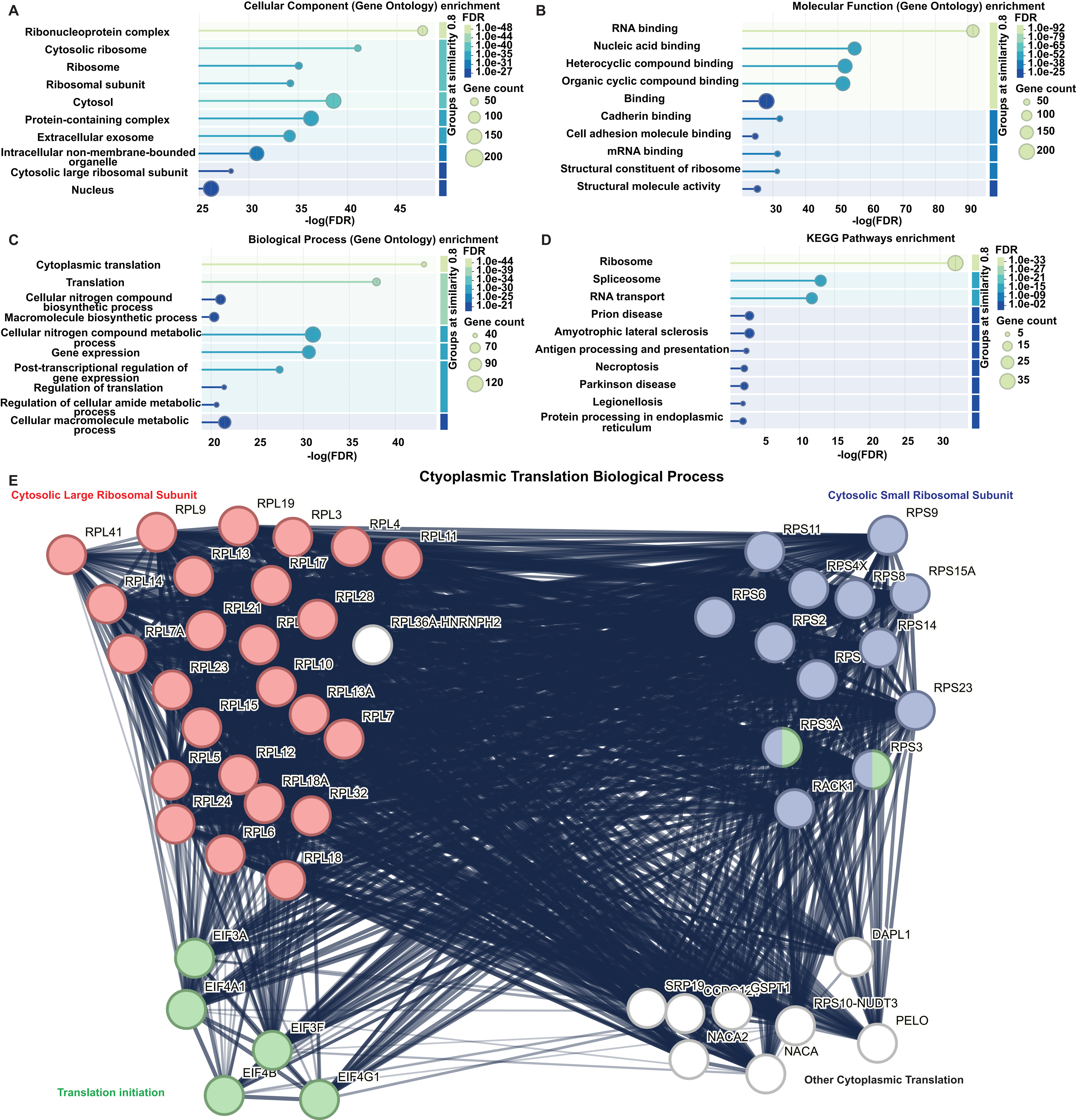

### Figure S2

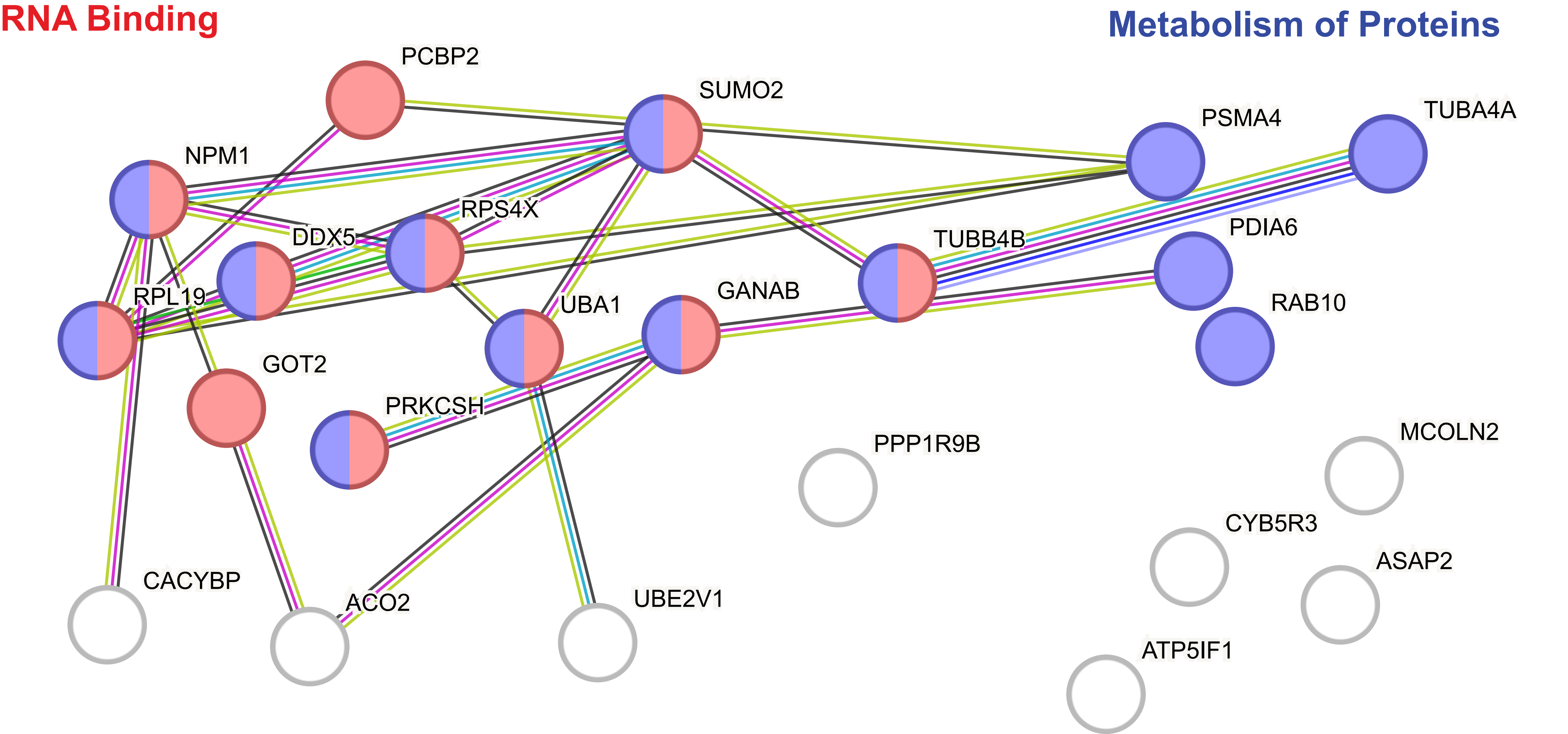

### Figure S3

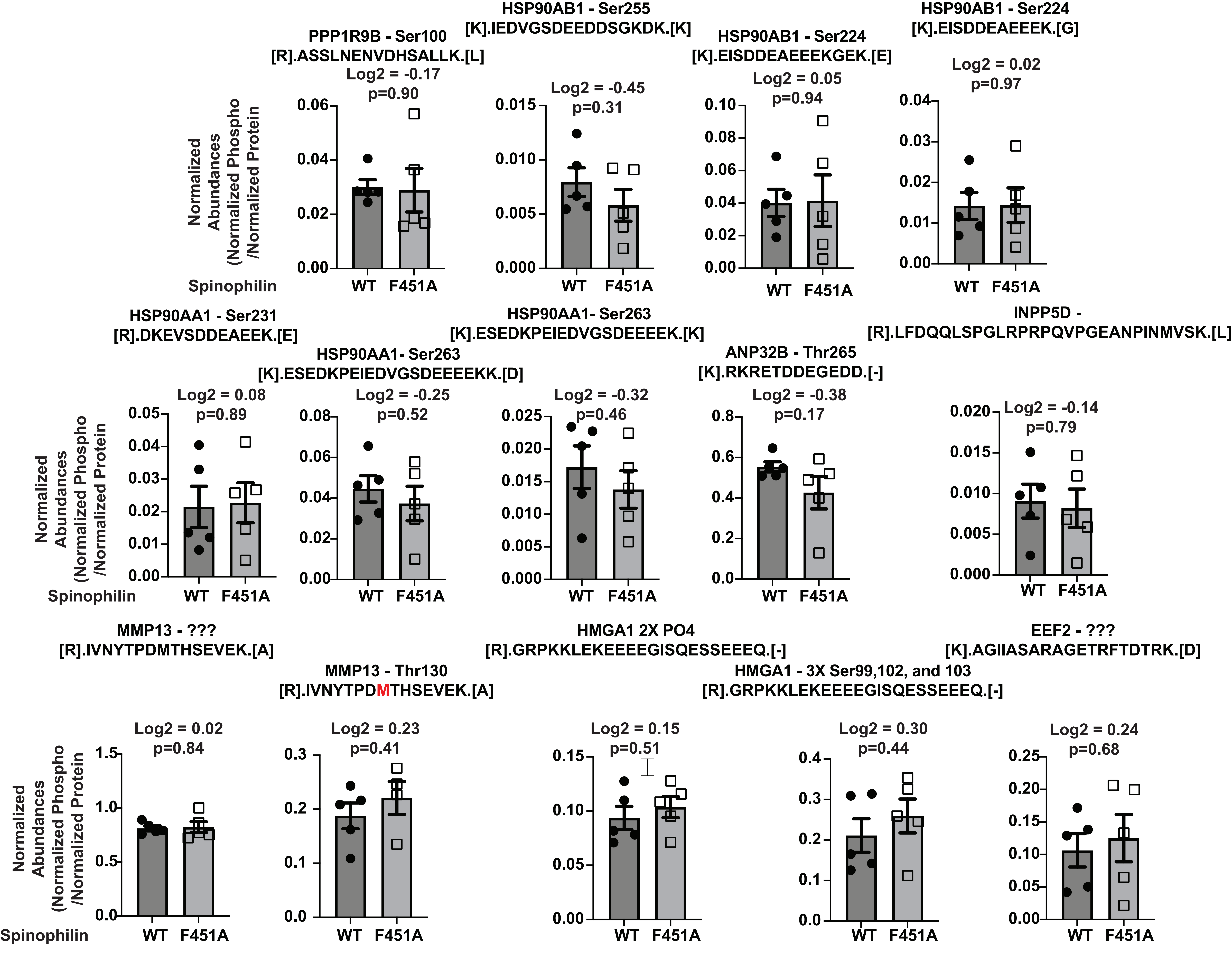

### Figure S4

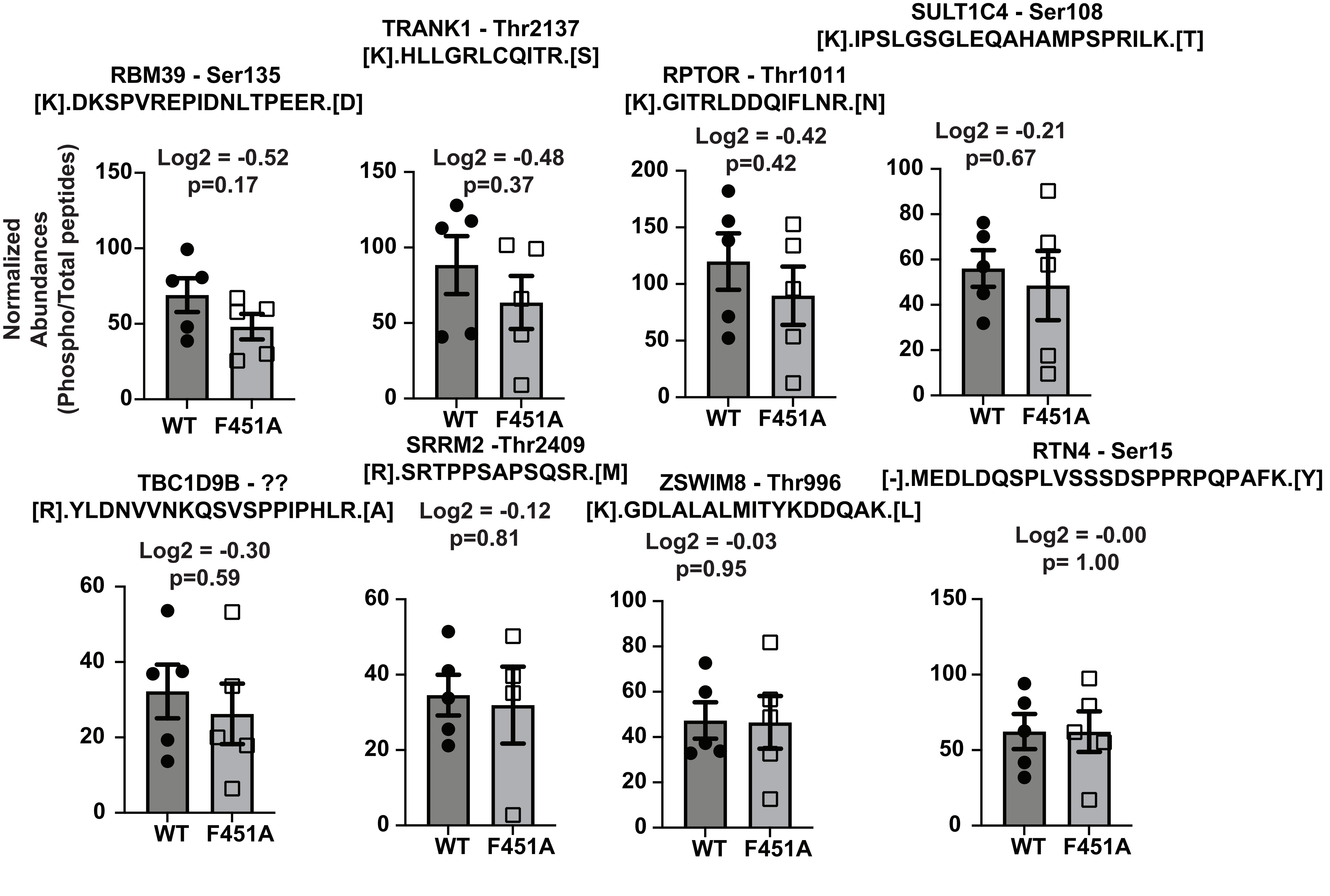

### Supplemental Westerns

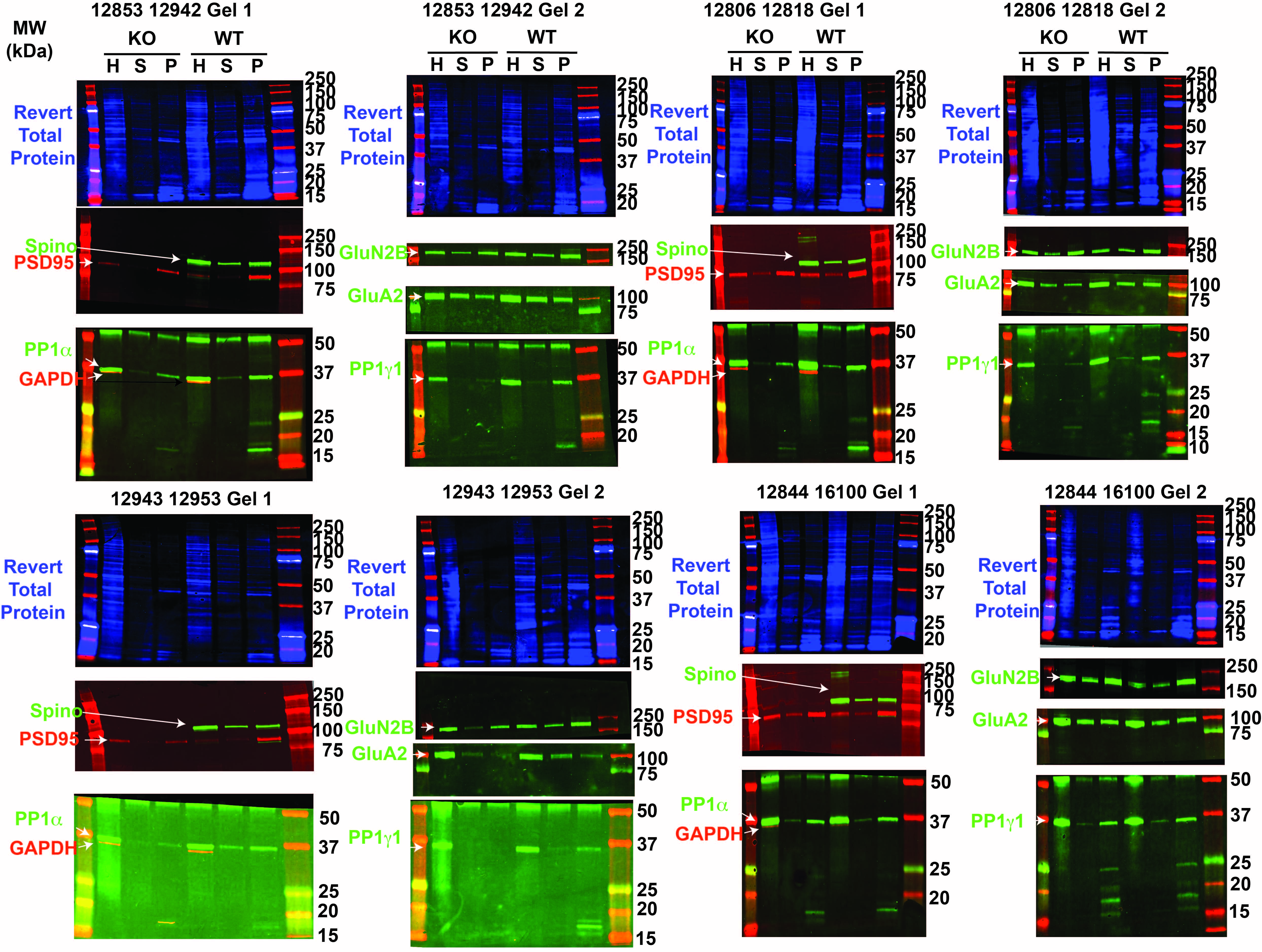
